# A Machine Learning Approach to Unmask Novel Gene Signatures and Prediction of Alzheimer’s Disease Within Different Brain Regions

**DOI:** 10.1101/2021.03.03.433689

**Authors:** Abhibhav Sharma, Pinki Dey

**Affiliations:** School of Computer and System Sciences, Jawaharlal Nehru University, New Delhi110067, India; School of Biotechnology and Biomolecular Sciences, University of New South Wales, Sydney 2033, Australia

**Keywords:** Alzheimer’s disease, Machine learning, Biomarkers, Gene expression, Feature Selection, Classification

## Abstract

Alzheimer’s disease (AD) is a progressive neurodegenerative disorder whose aetiology is currently unknown. Although numerous studies have attempted to identify the genetic risk factor(s) of AD, the interpretability and/or the prediction accuracies achieved by these studies remained unsatisfactory, reducing their clinical significance. Here, we employ the ensemble of random-forest and regularized regression model (LASSO) to the AD-associated microarray datasets from four brain regions - Prefrontal cortex, Middle temporal gyrus, Hippocampus, and Entorhinal cortex- to discover novel genetic biomarkers through a machine learning-based feature-selection classification scheme. The proposed scheme unrevealed the most optimum and biologically significant classifiers within each brain region, which achieved by far the highest prediction accuracy of AD in 5-fold cross-validation (99% average). Interestingly, along with the novel and prominent biomarkers including CORO1C, SLC25A46, RAE1, ANKIB1, CRLF3, PDYN, numerous non-coding RNA genes were also observed as discriminator, of which AK057435 and BC037880 are uncharacterized long non-coding RNA genes.

## 1. Introduction

Currently, 40-50 million people around the world are living with dementia and this number has doubled from 1990 to 2016^1^. Alzheimer’s disease being the most common form of dementia is expected to rise notoriously with the aging population. With the increase in its incidence, the expenses are also rising. It is estimated that in 2010 alone, Alzheimer’s disease had cost the world $604 billion^2^ and is expected to incur a global AD-associated healthcare cost of $2 trillion by 2030 affecting more than 131 million people by 2050^3^. Hence, Alzheimer’s disease is rapidly emerging as critical global health and economic challenge that has prompted vigorous scientific investigations to identify underlying genetic risk factors and regulatory markers, to suppress the estimated healthcare burden by early detection, especially at pre-symptomatic stages. Much research is performed on the late occurring hallmarks of AD^4–6^ such as neurofibrillary tangles, amyloid plaques, neuronal tangles, etc. Although these findings hold some important diagnostic values, the overall therapeutic contributions of these late occurring hallmarks of AD remain murky^4^. Moreover, clinical trials indicate that patients with AD show a varied pattern of symptoms and varying responses to a particular therapy that substantiates several pathological causes, making AD even more intricate to investigate^7^.

In recent years, data generated through high throughput gene expression profiling has opened new avenues for a better understanding of the complex disease mechanism and pathways at a molecular level^8, 9^. However, the huge dimension, low sample size, and noise in high-throughput gene expression data make it challenging to identify embedded patterns within the dataset. Currently, the methods to identify the most explaining gene subsets by data reduction and feature selection in the context of gene expression profile dataset analysis are broadly classified into two classes^10^: (i) marginal filtering method^11, 12^ and (ii) wrapper (embedded) method^13, 14^. The marginal filtering further is subdivided into two types namely, univariate and multivariate. Some examples of univariate filtering methods are paired t-test (TS), F-test (FT), and Pearson Correlation coefficient (PC)^11–13^. Some multivariate filtering approaches are Analysis of variance (ANOVA), F-score, feature selection based on correlation (CFS), and Max-Relevance-Max-Distance (MRMD)^15–18^. Using these methods, weights are assigned to the features (genes), and the genes with higher weights are considered to be the biologically important features. Although the filtering methods are computationally less expensive than the latter approach, they have significant shortcomings i.e. (i) most of the marginal filtering only accounts for the marginal contribution of a gene candidate while completely ignoring the interdependencies among the genes, and (ii) the absence of classification process. The filtering method doesn’t corroborate the classification accuracy of the selected features, reducing its clinical credibility^14^. However, the shortcomings of marginal filtering^19, 20^ can be overcome by wrapper methods. Wrapper methods are a hybrid of learning algorithms and classifiers that iteratively search for the optimum set of features by corroborating the classification accuracy of each chosen subset of candidate features^10^. Although the wrapper methods are very computationally intensive for high dimensional gene datasets, the classification accuracies obtained by the feature subsets identified by these methods are noticeably high^14^. In addition to this, machine learning models are empowered with efficient dimension reduction and feature selection methodologies to overcome the curse of dimensionality within the gene expression dataset^21^. Over time, many studies have employed machine learning models on microarray datasets to develop robust predictive models for identifying disease onset and prognosis of complex diseases such as cancer^22–25^.

Several studies have extensively leveraged machine learning models to identify biomarkers of AD from phenotypic data such as magnetic resonance imaging^26^. However, the identification of molecular signature underlying the mechanism of AD through gene expression profiles of demented patients remains largely unexplored^27^. In this direction, few studies have employed machine learning on gene expression data to delineate the potential differentially expressed genes (DEGs) within the AD-affected brain^28–31^. These studies have successfully used several state-of-the-art machine learning algorithms such as random forest, decision trees, support vector machines, and deep learning models to the feature selection and classification paradigm^32–35^. Although highly innovative, these methods had their own shortcomings such as, the proposed schemes within many of these methods were able to reduce the dimensions (number of features) but they remained mute on demonstrating the discriminative potential of the acquired DEGs, thus fails to vindicate the practical biological relevance of the obtained geneset. (ii) The majority of these studies incorporated only a small set of samples (usually <30), thus the results remained insufficiently descriptive and have low interpretability^32^.

Our objective here is to probe the difference in the gene expression levels within different brain regions of AD patients and non-demented controls, to identify the highly discriminating and biologically relevant gene signatures for AD through the wrapper (embedded) approach. We exclusively probe the Prefrontal cortex (PFC), Middle temporal gyrus (MTG), Hippocampus (H), and Entorhinal cortex (EC) as these regions are the most vulnerable to neurodegenerative diseases^36–38^. To retain the most significant and biologically relevant markers, we conceptualized a simple feature-selection and classification scheme based on the ensemble of random forest (RF) and regularized regression model; plugged with the best-configured classifier to obtain maximum classification accuracy in a 5-fold cross validation test (see **Fig 1)**. In addition to validating our finding by integrating biological knowledge through systematic literature review, GeneMania^39^, and STRING^40^ network analysis; we also corroborate the biological relevance of the obtained gene signatures by quantifying their disease discriminative power for the gene expression data obtained from the Visual Cortex (VC) and the Cerebellum (CR) of both AD affected and control brains. Through this work, we attempt to determine the signatures underlying AD and to formulate an efficient disease identification scheme whose clinical applications could further be extended for other diseases of altered expression.

**Fig 1.**
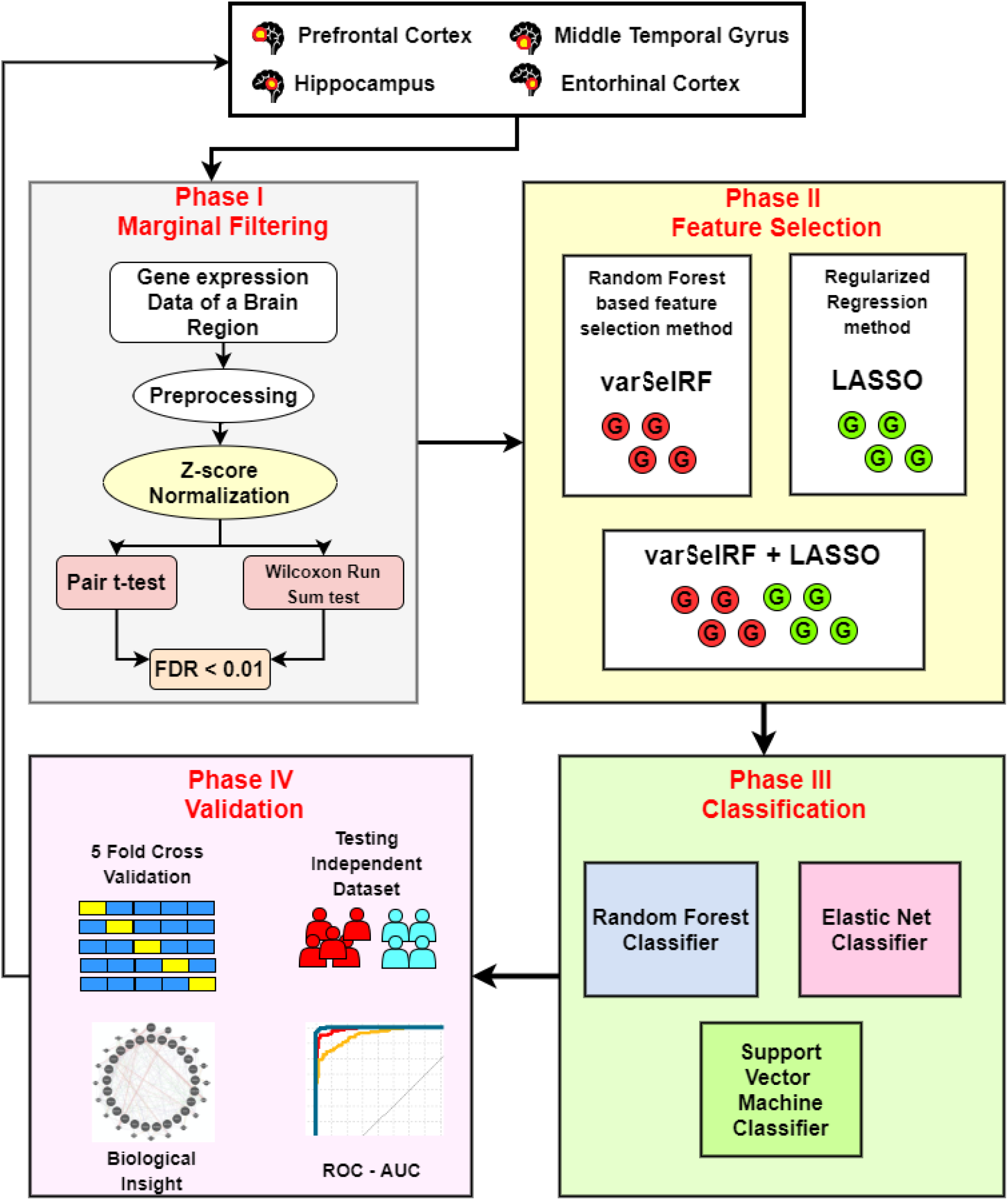
Schematic representation of the Machine Learning workflow to identify potential biomarkers for AD. The gene expression data for a given brain region is processed in the phase I. The features are then identified using wrapper methods (phase II). Subsequently in phase III and phase IV, the discriminative power and the biological relevance of the identified geneset is quantified and validated.

## 2. Materials and Methods

### 2.1 Dataset

We extracted the AD-associated gene expression datasets from the public functional genomics data repository NCBI-GEO database (http://www.ncbi.nlm.nih.gov/geo/). “Alzheimer’s” was used as a keyword to query all the experimental studies that have probed the gene expression profile within the brain tissues of AD patients against that of the non-demented healthy controls. The brain regions of our interest are the prefrontal cortex (PFC), middle temporal gyrus (MTG), hippocampus (H), and entorhinal cortex (EC). Datasets of only those studies were used that have performed microarray expression profiling and have a sample size of ≥15 for each type of brain tissue. This resulted in eight different studies, from which the samples of four brain tissue types (PFC, MTG, H and EC) were separated and grouped accordingly. This way we obtained a large sample size for each brain region. **Table 1** presents a summary of the expression datasets that are finally incorporated in this work. Each of these studies vary in terms of experimental design and measurements, that require special treatment to screen out definite AD and control samples for which we provided a detailed description of each dataset in supplementary **Table S1**.

**Table 1.**
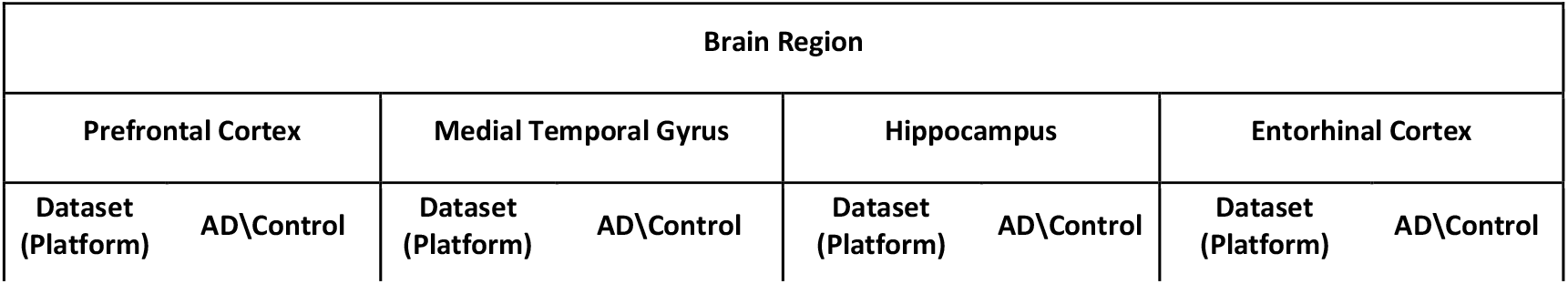
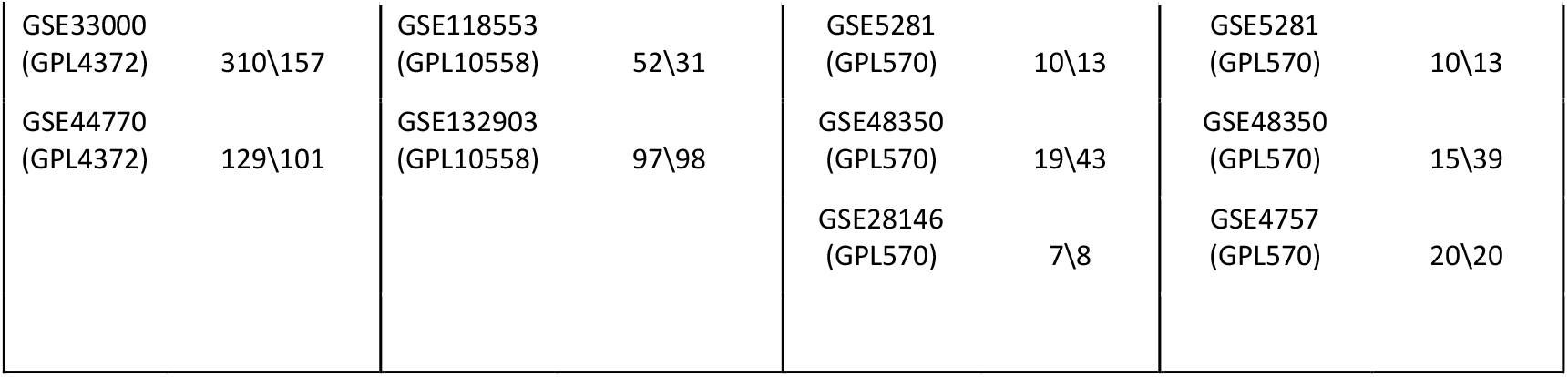
The gene expression datasets of Alzheimer’s Disease for four different brain regions.

The computation was carried out on an Intel (R) Core (TM) i5-4310U, 16 GB RAM, and 64-bit OS Win 10 configuration. Method implementation and experiments were conducted using R version 4.0.3. The schematic representation of machine learning workflow to identify potential biomarkers of AD is shown in **Fig 1**.

#### 2.1.1 Dataset integration and Pre-processing

To increase our sample size for statistically augmented results, we integrated at least two gene expression datasets for each brain region. However, the merging of the expression dataset is challenging because, (i) the platform over which the datasets were originated varies. Each type of platform measures the expression level of a particular set of genes which could be highly different from the gene repertoire of the other platforms; (ii) Due to adopting varying protocols, platforms and processes, different experiments contain various non-biological technical variations in the measurements^41^. These variations can induce a batch effect to the profiles that is potent to confound the true biological variations, thus may indicate misleading conclusions. To overcome these challenges, we essentially chose only those datasets to merge that were generated over a common platform. To subdue the batch effect, we standardized the expression profile of each sample, thus accounting for only the distribution of the gene expression^42^. For each dataset, the probe IDs were mapped to their respective Entrez gene IDs and Genbank Accession IDs that are annotated in the dataset’s corresponding platform table. In the case of duplicated gene IDs, the candidate with the maximum interquartile range was kept for further analysis. It was only after this step, we z-score normalized each sample to capture the distribution of the expression. We evaluated the p values for each gene candidate using both paired t-test and Mann Whitney U test, followed by its corresponding FDR correction for PFC and MTG due to their large sample size (> 200). Finally setting p<0.05 and FDR < 0.01, we prune our fully merged and pre-processed datasets for feature selection and classification.

### 2.2 Feature selection

As aforementioned, the merged gene expression datasets were the compilation of measurements from different samples but were generated from the same brain tissue, thus capturing the crucial biological basis for such expression within that particular brain region. To fetch the important independent players (gene candidates) underlying these expression levels, we employed two highly efficient feature selection methods; (i) Variable selection using Random forest method^43^ and (ii) Lasso regression method^44^. The parent models of these methods are probably the most pervasive machine learning algorithms i.e., random forest and generalized regression model respectively. The formalisms and the implementations of these methods are elaborated in the following sub sections.

#### 2.2.1 Variable Selection Using Random Forest (varSelRF)

The random forest algorithm developed by Breiman L.^45, 46^ uses the ensemble of regression trees for classification. Employing a bootstrap sample of the data, the classification tree is built. The candidate set of variables at each split of the tree is a random subset of the variables^44, 47^. In this way, RF incorporates bootstrap aggregation (bagging) and feature selection to build trees. To obtain low-bias trees, each tree is grown fully, and then bagging and random selection of variables is performed to facilitate low correlation of the individual trees^43^. For each fitted tree, RF registers a measure of error rate (OOB error) based on the out-of-bag cases (samples that have no contribution in the tree formation) that have very crucial applications in data reduction and feature selection. A detailed description of the algorithm underlying RF is provided in the supplementary text. Based on the characteristics of the RF algorithm, Ramón et al.^43^ formularized a feature selection model called varSelRF. This method is available as a package “*varSelRF*” on CRAN repository^48^. varSelRF iteratively fits random forests and selects a set of features (genes) that retains a minimum OOB error rate. Exploiting the embedded classification process, varSelRF returns a small subset of important genes while augmenting the predictive performance. This approach has already been incorporated in several literatures and has shown promising results^49–52^. The rationale to employ varSelRF in our framework is (i) the method returns a small set of gene candidates that has low correlation and high predictive power^52^ and (ii) RF based approach requires a less fine-tuning of parameters as the default parameter values often deliver the best performance^53^.

#### 2.2.2 Regularized regression models

*Least Absolute Shrinkage and Selection Operator* (LASSO) is a type of regularization regression method to fit a generalized linear model. Based on the idea of penalizing the regression model (L1-norm), LASSO squashes the regression coefficient to zero for the variable that has the least contribution to the model. This way the LASSO regression model has an optimal feature selection capability. LASSO regression is an alternative regression approach to Ridge regression that too is based on penalizing the model but follows a L2-norm^44^.

For a given population *X*, let *x*_*ij*_ be the *i*^*th*^(1 ≤ *i* ≤ *n*) observation of the *j*^*th*^(1 ≤ *i* ≤ *p*) variable and let *y*_*i*_ be the corresponding label of the *i*^*th*^ instance. For each *p* variable, the regularized regression model estimates the regression coefficient *β*_*j*_(1 ≤ *i* ≤ *p*) by minimizing the sum of squared error (eq. 1) along with a constraint on the coefficients ∑ *J*(*β*_*j*_) ≤ *t*^44, 54, 55^.

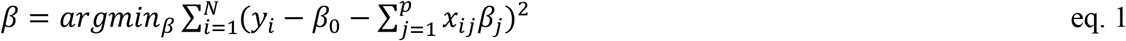

For LASSO the coefficient *J*(*β*_*j*_) = |*β*_*j*_| and in the Ridge *J*(*β*_*j*_) = *β*_*j*_^2 42, 51^. This way LASSO regression tranculates the coefficient of the non-contributing variable to zero while Ridge shrinks the coefficient close to zero, delineating LASSO as an efficient feature selection model. The LASSO obtains the *β*_*j*_ estimate by minimizing eq. 2.

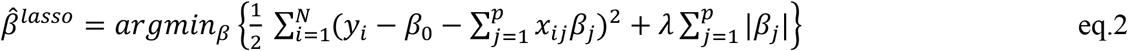

where λ is the penalty parameter that determines the shrinkage proportion and is often determined using cross-validation^44^. LASSO retains an excellent performance for the situations when (i) the data has very high dimension and low sample and (ii) few variables explain the majority of data (have large coefficient) and the remaining variable has very low predictive potentials^44^. Moreover, LASSO has some significant advantages such as (i) LASSO efficiently handles the multicollinearity within the features and returns highly independent features and (ii) Being computationally less expensive, LASSO retains the optimal gene candidates faster. These characteristics of LASSO befit the gene expression data as a feature selection model. LASSO has elucidated excellent performance in numerous studies^55–58^, delineating as a very promising feature selection model. The variables with relative scaled importance >10 was considered significantly important.

However, studies have indicated that L1-norm (LASSO) is not universally dominant over the L2 norm (Ridge). However, to improve computational tractability Zou et al.^59^ proposed a relatively new penalty called the Elastic Net, built as an intelligent compromise between LASSO and Ridge penalty. For Elastic Net, the *J*(*β*_*j*_) (coefficient constrain) is:

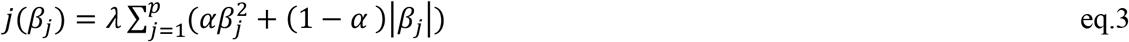

where the new α constant is introduced that regulates the intensity of LASSO and Ridge penalties. Elastic Net handles multicollinearity more efficiently than LASSO by accounting for every correlated pair during training^44, 59^. Elastic Net has better performance on many occasions, however, there are only a few studies that corroborate the same^60^. Although we have employed LASSO as a feature selection, we leverage the Elastic Net classifier to test the determinative power of all the selected features combined due to its high efficiency towards multicollinearity. The R package “caret” was used to implement LASSO and Elastic Net^61^.

#### 2.2.3 Multiplicity Problem

For microarray dataset, the problem of multiplicity can cast a false sense of trust in the genes identified by wrapper approach. The basis of multiplicity problem has been explained in detail in the supplementary text. Subscribing to the notion of the studies investigating the problem of multiplicity^62–65^, we lend credence to the combined set of genes that were obtained by both the methods (varSelRF and LASSO); and exclusively probed the biological significance of the common and repeatedly selected gene candidates.

### 2.3 Classification model

Classification modeling led by feature selection is a crucial phase of the paradigm, that depicts the clinical application of the selected gene candidates. Although the embedded classifier within the wrapper method leverages the classification accuracy to quantify the importance of a gene subset, but in the context of therapeutic application it is very crucial to corroborate the best suiting classification model that improves the prediction accuracy. In this work, we employed Support Vector Machines (SVM), Random forest classifier, and Elastic Net classifier and performed a comparison study. We also probed the classification efficacy of these models for the gene candidates obtained by (i) varSelRF, (ii) LASSO and (iii) combined gene subsets retained by both varSelRF and LASSO. An overview of RF and SVM classifiers is provided in the supplementary text. The R package “random Forest” and “e1071” were used to implement the RF^53^ and SVM^66^ respectively.

### 2.4 Assessment

#### 2.4.1 Model Assessment

We assess the prediction power of the selected gene candidates through SVM, RF, and Elastic Net classifiers. Exploiting the relatively large sample size due to the merging of gene datasets, we perform a 5-fold cross validation method to judge the external prediction power of the gene set as well as of the classification model with a high level of certainty. To compare the efficacy of the models we measure the following metrics.

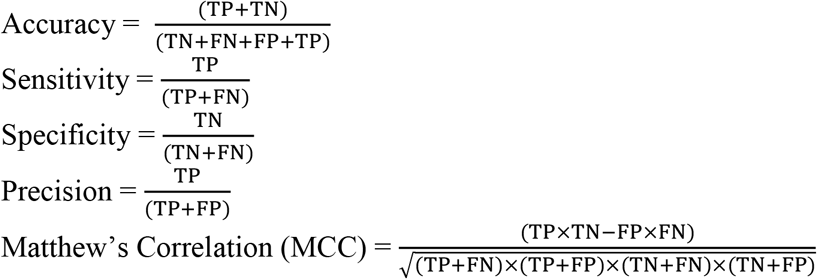

Here TP, TN, FN and FP represent true positive, true negative, false negative and false positive predictions respectively made by the classification model for each brain region (AD state is denoted positive and non-demented healthy state is denoted negative). For further comparative analysis, we plot the receiver operating characteristics (ROC) curve and compared the area under the curve (AUC) obtained by the model for different brain tissue.

#### 2.4.2 Feature Assessment

We assess the biological significance of the feature set obtained by our framework by integrating the biological knowledge through a systematic literature review. We have used GeneMania^39^ and STRING^40^ network analysis to identify the co-expression, genetic and physical interactions among the obtained biomarkers of AD and also with the previously well-known AD genes. Using the same, we also delineated the networks (hub genes) associated with our obtained molecular signatures to deliver deeper insight into the mechanism of AD in different brain tissues. To further corroborate the biological meaningfulness of the obtained markers, we tested the discriminative power of these markers to classify AD patients from non-demented controls for two different brain regions, Visual Cortex (VC) and the Cerebellum (CR). This way, not only the biological relevance is unmasked quantitatively, the therapeutic application of the proposed framework is also depicted.

## 3. Results

After marginal filtering in the first phase (Phase I), we obtained 26,593, 13,037, 3,268 and 10,029 genes for PFC, MTG, H and EC respectively, that was further processed to identify DEGs using the varSelRF and LASSO method (Phase II). In the following section, we shed light on the obtained features as well as their discriminative power when treated with different benchmark classifiers.

### 3.1 The Obtained Features

For each brain region, we employed varSelRF and LASSO on the gene candidates from phase I. varSelRF is based on RF that has the inner nature of being purely random and performs random sampling within the algorithm. This leads to slightly varying results when implemented multiple times. Therefore, for each brain region, we implemented varSelRF for five times and considered each selected candidate as important feature. The tuned hyperparameter sets for varSelRF and LASSO are provided in the supplementary **Table S2**. The features obtained are summarized in **Table 2**. We found that LASSO obtained a higher number of candidates than varSelRF for the brain region with a large sample size and vice versa.

**Table 2.**
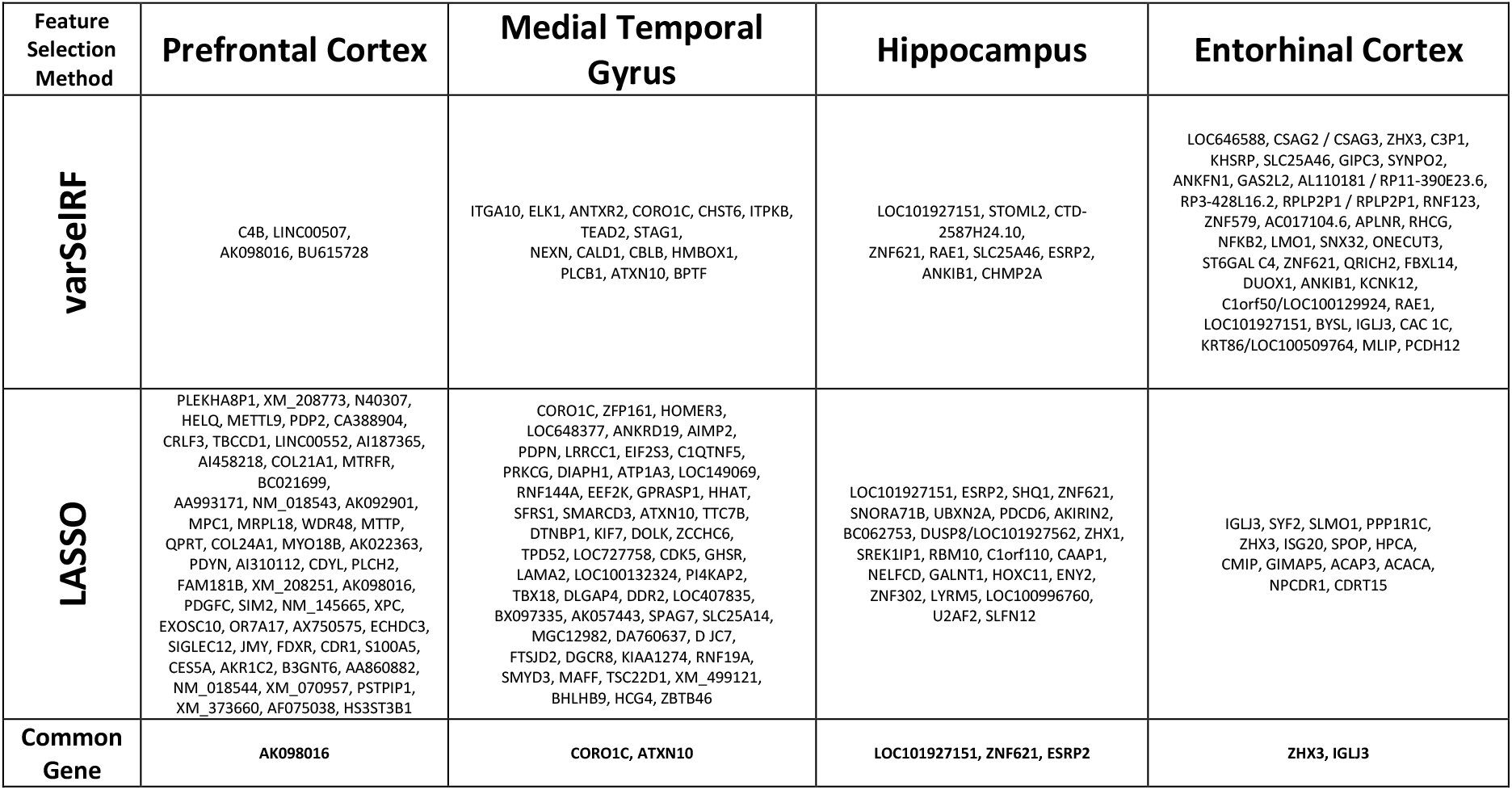
The gene biomarkers obtained for different brain regions using varSelRF and LASSO methods.

Both the models largely identified a varying set of markers; however few gene candidates were commonly identified by both methods. Of interest, the majority of these commonly identified markers are closely associated with neurodegenerative disorders, depicting the biological significance of the models. In addition to the common genes identified by the models, there were common regulatory gene candidates within the brain regions **(see Fig S1)**. The common biomarkers found within the H and EC region are ZNF621, SLC25A46, RAE1, and ANKIB1. Among these biomarkers, RAE1, ANKIB1, and SLC25A46 have been reported to be prominently involved in several neurodegenerative disorders. The RAE1 protein is found to be the interacting partner of Huntingtin protein aggregates^67^ and experimental evidence of early ageing associated phenotypes is reported in Rae1 haplo-insufficient mice^68^. ANKIB1 is also found to be associated with Cerebral cavernous malformations^69^. Another potential biomarker that has been associated with neurodegenerative disorders is SLC25A46. A study by Abram’s et al. has experimentally shown that the mutations in the SLC25A46 genes can lead to the degeneration of optic and peripheral nerve fibers^70^. Also, loss of function in the SLC25A46 gene leads to lethal congenital and peripheral neuropathy^71, 72^. Although these genes have been extensively studied for different neurological disorders, their role in Alzheimer’s disease is yet to be exclusively explored. Our models were also able to unravel the participation of non-coding RNAs, identifying 9 non-coding RNAs within the brain regions. Among the non-coding RNAs, we found two long non-coding RNAs, AK057435 and BC037880 in the prefrontal cortex and the hippocampus region respectively that are classified as potential biomarkers. Since long non-coding RNAs are known to play an important role in human neurological development and cognition, experimental characterization of these biomarkers can help to elucidate the role of long non-coding RNAs in Alzheimer’s disease.

### 3.2 Classification

To determine the classification potential of the obtained gene set for each brain region, we built three benchmark classification models (SVM, random forest and Elastic Net). Performing extensive machine learning experiments, we made an attempt to identify the best pair of feature-selection and classification models in the context of disease class prediction. For each of the four brain regions, we applied three different best-configured classification model to the gene set obtained through varSelRF, LASSO and finally to the combine pool of gene set (varSelRF + LASSO), depicting a total of 9 scenarios to identify the best performing combination. The classification performance was assessed through a 5-fold cross validation method. **Table 3** represents a complete summary of the assessment metrics obtained for each possible scenario. In our study, the proposed framework has obtained foremost the highest AD prediction accuracy than any previous studies in a similar paradigm to our knowledge to date. For the prefrontal cortex and hippocampus, the scheme has even obtained 100% prediction accuracy.

**Table 3.** Performance comparison of the three different classification models (SVM, RF, Elastic Net) applied to the gene set obtained through varSelRF, LASSO and varSelRF + LASSO for the four brain regions, namely Prefrontal cortex, Middle Temporal Gyrus, Hippocampus and Entorhinal Cortex.

| Feature Selection Method | Model | Brain Region |  |  |  |  |  |  |  |  |  |  |  |  |  |  |  |  |  |  |  |
| --- | --- | --- | --- | --- | --- | --- | --- | --- | --- | --- | --- | --- | --- | --- | --- | --- | --- | --- | --- | --- | --- |
|  |  | Prefrontal cortex |  |  |  |  | Middle temporal gyrus |  |  |  |  | Hippocampus |  |  |  |  | Entorhinal cortex |  |  |  |  |
|  |  | Acc | Sen | Spe | Pre | Mcc | Acc | Sen | Spe | Pre | Mcc | Acc | Sen | Spe | Pre | Mcc | Acc | Sen | Spe | Pre | Mcc |
| varSelRF | SVM | 0.93 | 0.95 | 0.90 | 0.91 | 0.85 | 0.86 | 0.85 | 0.87 | 0.89 | 0.73 | 0.90 | 1.00 | 0.80 | 0.84 | 0.82 | 0.91 | 0.95 | 0.80 | 0.80 | 0.75 |
|  | RF | **0.95** | **0.95** | **0.94** | **0.94** | **0.89** | 0.87 | 0.88 | 0.87 | 0.88 | 0.74 | **0.94** | **1.00** | **0.88** | **0.91** | **0.89** | **0.95** | **1.00** | **0.85** | **0.87** | **0.84** |
|  | ElasticN | 0.93 | 0.97 | 0.90 | 0.90 | 0.87 | **0.88** | **0.86** | **0.89** | **0.90** | **0.75** | 0.93 | 0.97 | 0.88 | 0.91 | 0.87 | 0.94 | 1.00 | 0.82 | 0.84 | 0.81 |
| LASSO | SVM | 0.99 | 1.00 | 0.99 | 0.99 | 0.99 | 0.96 | 0.96 | 0.95 | 0.96 | 0.91 | 0.79 | 0.81 | 0.77 | 0.78 | 0.59 | 0.91 | 0.92 | 0.90 | 0.95 | 0.79 |
|  | RF | 0.97 | 0.97 | 0.96 | 0.96 | 0.93 | 0.91 | 0.91 | 0.91 | 0.92 | 0.81 | 0.83 | 0.83 | 0.83 | 0.84 | 0.66 | 0.92 | 0.91 | 0.92 | 0.88 | 0.78 |
|  | ElasticN | **1.00** | **1.00** | **1.00** | **1.00** | **1.00** | **0.99** | **0.99** | **0.99** | **0.99** | **0.99** | **0.86** | **0.86** | **0.87** | **0.87** | **0.74** | **0.97** | **0.98** | **0.97** | **0.97** | **0.93** |
| varSelRF + LASSO | SVM | 0.99 | 1.00 | 0.99 | 0.99 | 0.98 | 0.95 | 0.95 | 0.95 | 0.96 | 0.91 | 0.99 | 1.00 | 0.97 | 0.98 | 0.97 | 0.92 | 0.95 | 0.82 | 0.84 | 0.78 |
|  | RF | 0.97 | 0.98 | 0.96 | 0.96 | 0.94 | 0.88 | 0.87 | 0.88 | 0.90 | 0.75 | 0.94 | 0.98 | 0.91 | 0.93 | 0.90 | **0.95** | **1.00** | **0.85** | **0.87** | **0.84** |
|  | ElasticN | **1.00** | **1.00** | **1.00** | **1.00** | **1.00** | **0.98** | **0.99** | **0.98** | **0.98** | **0.96** | **1.00** | **1.00** | **1.00** | **1.00** | **1.00** | 0.94 | 1.00 | 0.82 | 0.84 | 0.81 |

### 3.3 Performance Evaluation

It was observed that in the majority of the scenarios, the Elastic Net classifier obtained excellent performance, followed by the random forest classifier, while SVM performance remained low (**Fig 2)**. Substantiating the parent algorithms, both RF and Elastic Net classifier has performed higher for the gene sets obtain through their respective allied feature selection model i.e., varSelRF and LASSO respectively. Considering the problem of multiplicity, we substantiate the combined gene markers of varSelRF and LASSO over the gene set obtained by these individual methods. The ROC-AUC plot elucidates the superiority of Elastic Net over RF and SVM for three brain regions (PFC, MTG and H) while remaining slightly lower but highly competitive for the EC region (**Fig 3)**. The one explanation of low performance of Elastic Net for EC region is possible due to the very small sample to gene ratio.

**Fig 2.**
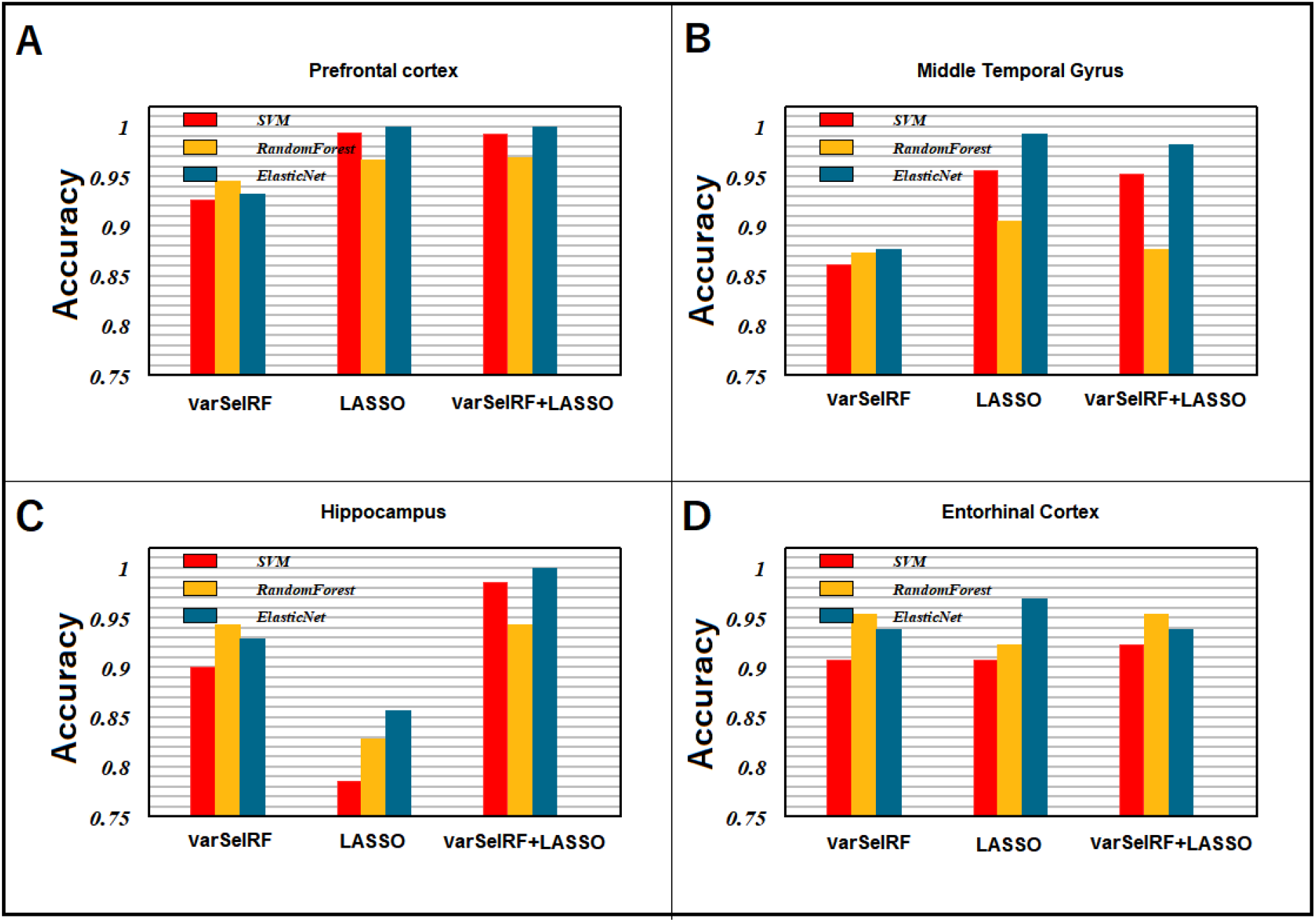
Prediction accuracy obtained by the SVM, Random Forest, Elastic Net classifier employed in the varSelRF, LASSO and varSelRF + LASSO for **(A)** Prefrontal cortex, **(B)** Middle Temporal Gyrus, **(C)** Hippocampus and **(D)** Entorhinal Cortex. The Elastic Net classifier obtained excellent performance in the majority of scenarios, followed by the random forest classifier and SVM. Genes obtained through LASSO with Elastic net classifier performed higher in PFC, MTG and EC region.

**Fig 3.**
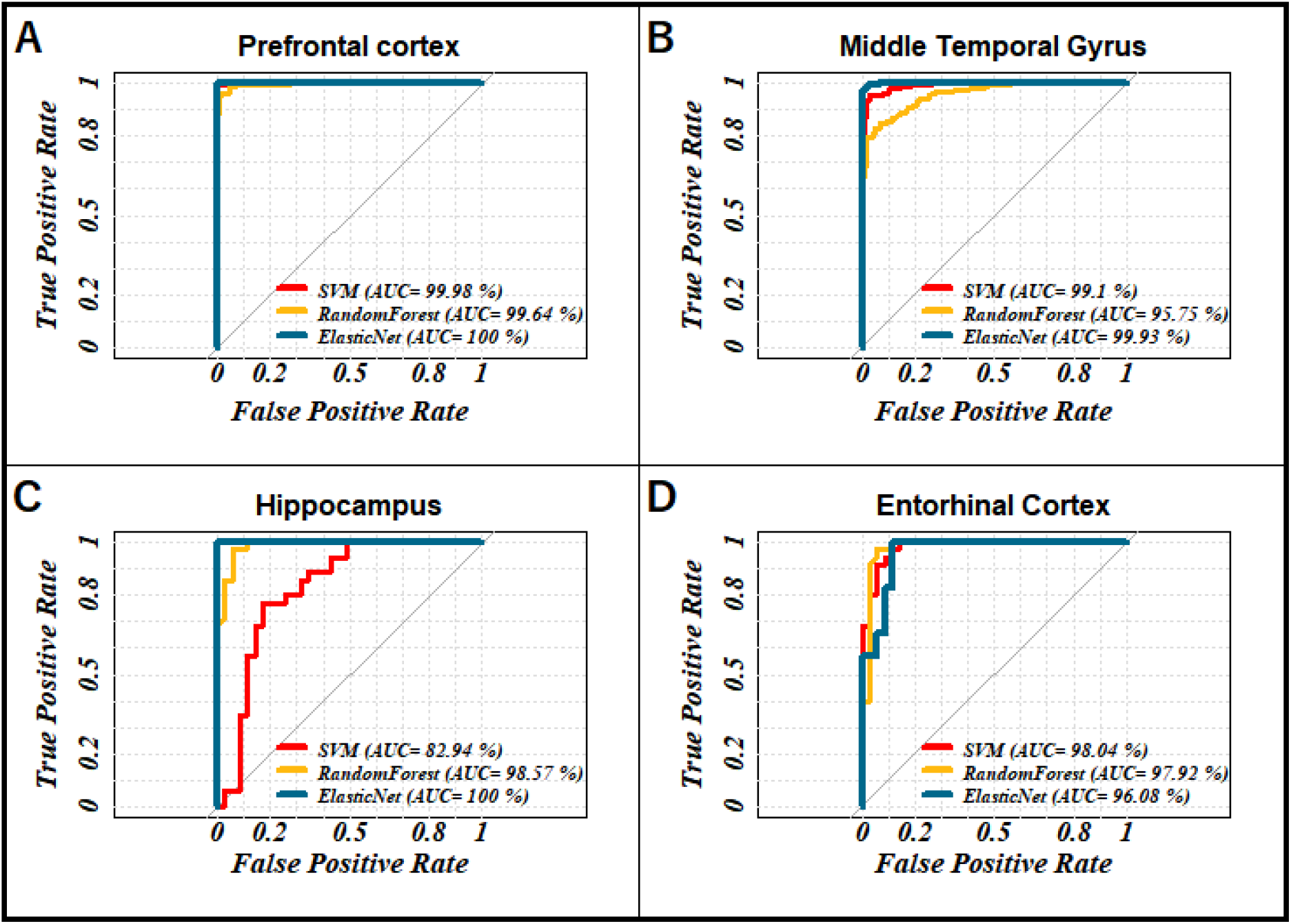
The classification performances to discover potential biomarkers in four brain regions. The ROC-AUC curves of Elastic Net, Random Forest and SVM classifiers for **(A)** Prefrontal cortex, **(B)** Middle Temporal Gyrus, **(C)** Hippocampus and **(D)** Entorhinal Cortex.

In addition to adopting a 5-fold cross validation method, we also took several other measures to establish the biological credibility of the identified gene candidates. We hypothesize that the gene markers obtained for one brain region hold some biological relevance for the adjacent brain region. We therefore evaluated the AD prediction potential of the gene subset of PFC (60 genes) for the gene expression data obtained from Virtual Cortex (VC) and Cerebellum (CR). VC and CR data were extracted from GEO NCBI database (GSE44771 and GSE44768). The sample size for VC and CR are both 230 with AD to control ratio of 129:101. We employed LASSO feature-selection only for the expression level of those 60 gene candidates that were identified as PFC markers on the VC and CR datasets. We find that the biomarkers of PFC displayed an excellent AD classification performance (5-fold CV) of 92% and 91% on VC and CR datasets respectively (see supplementary **Table S3**). The complete assessment metric obtained for VC and CR is provided in the supplementary **Table S4.** This quantitatively validates the biological meaningfulness of gene candidates obtained in our study.

## 4. Discussion

The formalism of the proposed framework has two integrated components (i) Identification of the AD associated crucial gene markers within each brain region and (ii) the disease class prediction. After carrying out an extensive comparative analysis and corroborating the problem of multiplicity, it is apparent that Elastic Net classifier has a remarkable potential for disease prediction when employed over the gene subset identified by multiple varieties of gene selection models (LASSO and varSelRF in this case). In addition to having outstanding AD predictive potentials, the markers identified through this framework are of high calibre in terms of explaining the expression level and multicollinearity.

**Fig 4A** illustrates the correlation heatmap for the expression level of the biomarkers obtained by LASSO and varSelRF for each brain region. We see that the biomarkers elucidated very low correlation, thus together they are of great relevance in the context of depicting the biological basis for the observed expression level. Although both feature selection models are immune to multicollinearity, the LASSO obtained significantly lower correlated markers than that of varSelRF, especially for the EC region. This is also apparent in the correlation density plot for the regions, where the density remained high near the centre for the geneset obtained through LASSO, while it remains inflated on the tails for the varSelRF obtained geneset (**Fig 4B**).

**Fig 4.**
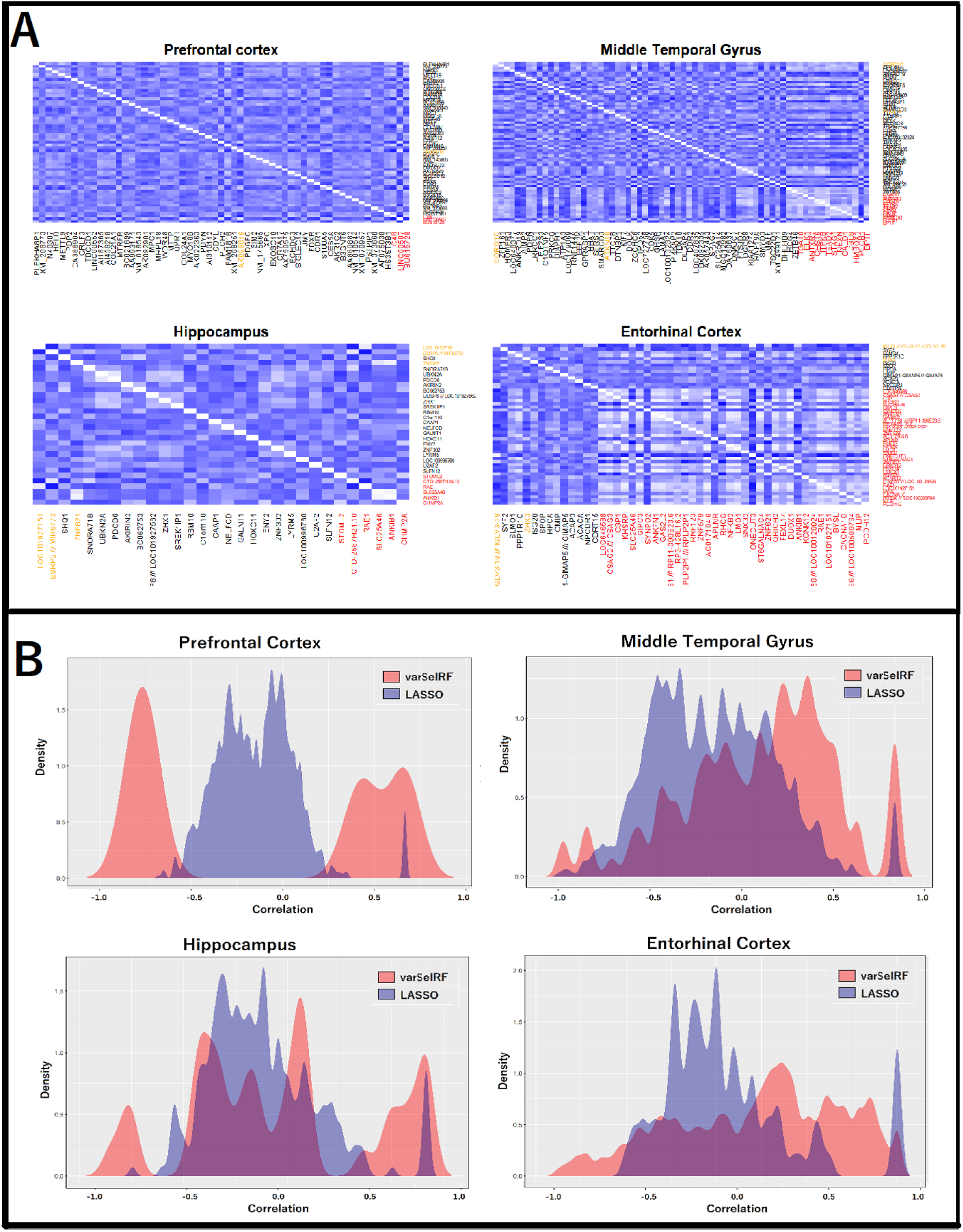
**(A)** The correlation heatmap (n x n, where n is number of biomarkers) for the expression level of the biomarkers obtained by LASSO and varSelRF method for each brain region. Every block in a heatmap plot represents correlation between the gene on each axis. Correlation ranges from −1 to +1. The shade corresponding to the values closer to zero indicate low linear trend between the two markers. The red labelled markers are the one that are obtained by varSelRF. The black labelled markers are the one that are obtained by LASSO. The orange labelled are the markers that were identified by both the models. **(B)** The density plot for the correlation values among the gene subset obtained by each type of feature selection model within different brain region. Density plot of correlation value for the markers obtained through varSelRF is shown in red. Density plot of correlation value for the markers obtained through LASSO is shown in blue. The density for the correlation value near to zero remained higher for LASSO comparative to varSelRF in every brain region.

### 4.1 Biological Insight

We performed a combination of biological network analysis and a comprehensive literature review to validate the biomarkers obtained in our study. We started with bioinformatics analysis of all the biomarkers obtained from our models and are listed in supplementary **Table S5**. We find the presence of potential biomarkers in all the chromosomes, except Chromosome Y. This may point towards the higher prevalence of AD in woman than in man^73^. The Chromosomes 1, 6, 17, 19 are found to contain the maximum number of biomarkers (**Fig S2**). Although most of the genes that are classified as biomarkers in our study are protein coding genes, some non-coding genes, such as LINC00552, LINC00507, MGC12982, HCG4, LOC101927151, NPCDR1, LOC646588 are also found to be the biomarkers of AD. These non-coding genes are novel and mostly uncharacterised.

Moreover, we identified 7 up-regulated and 6 down-regulated genes in the AD samples with respect to the normal ones by employing the GSE5281 expression data due to the availability of raw count. We considered p < 0.01 and |log2FC| ≥ 0.6 (FC, fold change) as cut-off criterion on different samples of H and EC brain regions from the GSE5281 dataset. Using this information, we identified the biomarkers that are up and down regulated (**Fig S3**). We find that some of the biomarkers are significantly downregulated in AD such as MLIP and STOML2. While the down regulation of STOML2 gene has been reported previously in AD patient’s samples^74^, the EC biomarker, MLIP can be clinically tested as a novel possible biomarker of AD.

We also performed GeneMania network analysis for all the biomarkers of each brain region (**Fig. S4–7**) and found that the biomarkers are not only co-expressed but share both physical and genetic interactions. Some of the highly interacting genes in the PFC are ECHDC3, PDGFC, MPC1, CRLF3, CDYL, FDXR that are also found to be co-expressed (**Fig S4**). Among these genes, the expression of ECHDC3 is found to be significantly higher in AD patients than non-AD patients from genome-wide association studies of more than 200,000 individuals^75^. Also, CRLF3 has been studied in neuronal aging rates in human brain regions^76^. In the EC region, we see that the biomarkers interact with each other by largely physical and genetic interactions **(Fig S5)**. In particular, ZNF621 and ISG20 are found to genetically interact with many of the other biomarkers. The ZNF621 gene has been recently reported as an upregulated gene in AD patients^77^. From the network analysis of the H region, we find extensive interactions of the biomarkers with each other, where the biomarkers not only are involved in physical and genetic interactions as well as co-expression and co-localization (**Fig S6**). Some of the highly interacting biomarkers of the H region are RBM10, SLC25A46, STOML2. It is interesting to note that both the SLC25A46, STOML2 protein are involved in mitochondrial dynamics and it has been proposed that mitochondrial dysfunction due to oxidative stress may be one of the earliest and prominent features of AD; and it has been experimentally shown that slower mitochondrial dynamics is correlated with reduced expression of STOML2 and MFN2^74^. The network analysis of MTG region shows that most of the biomarkers in the region genetically interact with each other, however, co-expression is also seen for some of the biomarkers such as CALD1, DNAJC7, TSC22D1, CMTR1, CORO1C (**Fig S7**). TSC22D1 is one of the most studied transcription factors that has also been reported as the potential new target for treating AD^78^. Hence, the biomarkers found by our models have not only been studied for different neuropathies but some of them are also reported as potential targets against AD. Also, we see that our biomarkers extensively interact with each other and thus, careful targeting of a potential biomarker can also help to regulate the biological functions of other biomarkers involved in various neuropathies.

### 4.2 Relationship between the biomarkers and AD genes

The most well-known genes that have the largest effect on the risk of developing AD are APOE, APP, PSEN1, and PSEN2^79^. Although we have not identified these genes in our study, the relationship between these AD genes and our biomarkers is worth analysing. To seek the potential interactions between the biomarker genes and the AD genes according to different brain regions, the STRING^40^ (*Search Tool for the Retrieval of Interacting Genes/Proteins*) tool was employed. Active interaction sources such as experimental data, public databases, text mining, computational prediction methods, and species limited to “*Homo sapiens*” are applied to construct the protein-protein interaction (PPI) networks. From the interaction networks shown in **Fig 5**, we see that the biomarkers of all the brain regions, except the hippocampus have interactions with the AD genes. In the prefrontal cortex, the biomarkers showing significant interactions with the AD genes are C4A, SIM2 and PDYN (**Fig 5A**). The complement pathway protein, C4A is found to be present in higher levels in patients with AD and represents the inflammation generally associated with neurodegenerative diseases^80^. The biomarker SIM2 is also supposed to serve as a noble target for Down’s Syndrome-related AD^81^. Although the PDYN gene is extensively studied in Huntington’s Disease^82^, its role in AD is yet to be explored. The interacting biomarkers with AD genes in the MTG region are CDK5, GHSR, PLCB1, ITPKB, HOMER3 (**Fig 5B**). CDK5 is gradually emerging as an obvious therapeutic target for AD because Cdk5/p25 is involved in two most important pathological hallmarks of AD, the formation of Aβ plaques and NFTs^83^. Also, in the current scenario, we see GHSR, PLCB1, ITPKB genes are considered to be promising therapeutic targets for AD^84–87^. Similarly, in the EC region, the interacting biomarkers are NFKB2, CACNA1C, APLNR (**Fig 5D**). The transcription factor NFKB2 has emerged as a potential target for AD prevention by targeted anti-inflammatory treatment to increase the time of disease onset^88^. Moreover, by targeting the calcium voltage-gated channel subunit alpha-1 C gene, CACNA1C by miRNA, studies have reported the inhibition of tau protein hyperphosphorylation in AD^89^. The apelin receptor protein, APLNR is also been recently studied as a potential target for several neurodegenerative diseases including AD as expression level alterations in apelin significantly affects the neuronal structure, calcium signalling, apoptosis, and autophagy etc^90^. From the analysis, we see that some of our biomarkers that closely interact with the well-known AD genes are also closely associated with various neurological disorders including AD. Future work requires the experimental testing of these gene biomarkers found in our study to identify the potential signature biomarker for efficient early diagnosis and treatment of AD.

**Fig 5.**
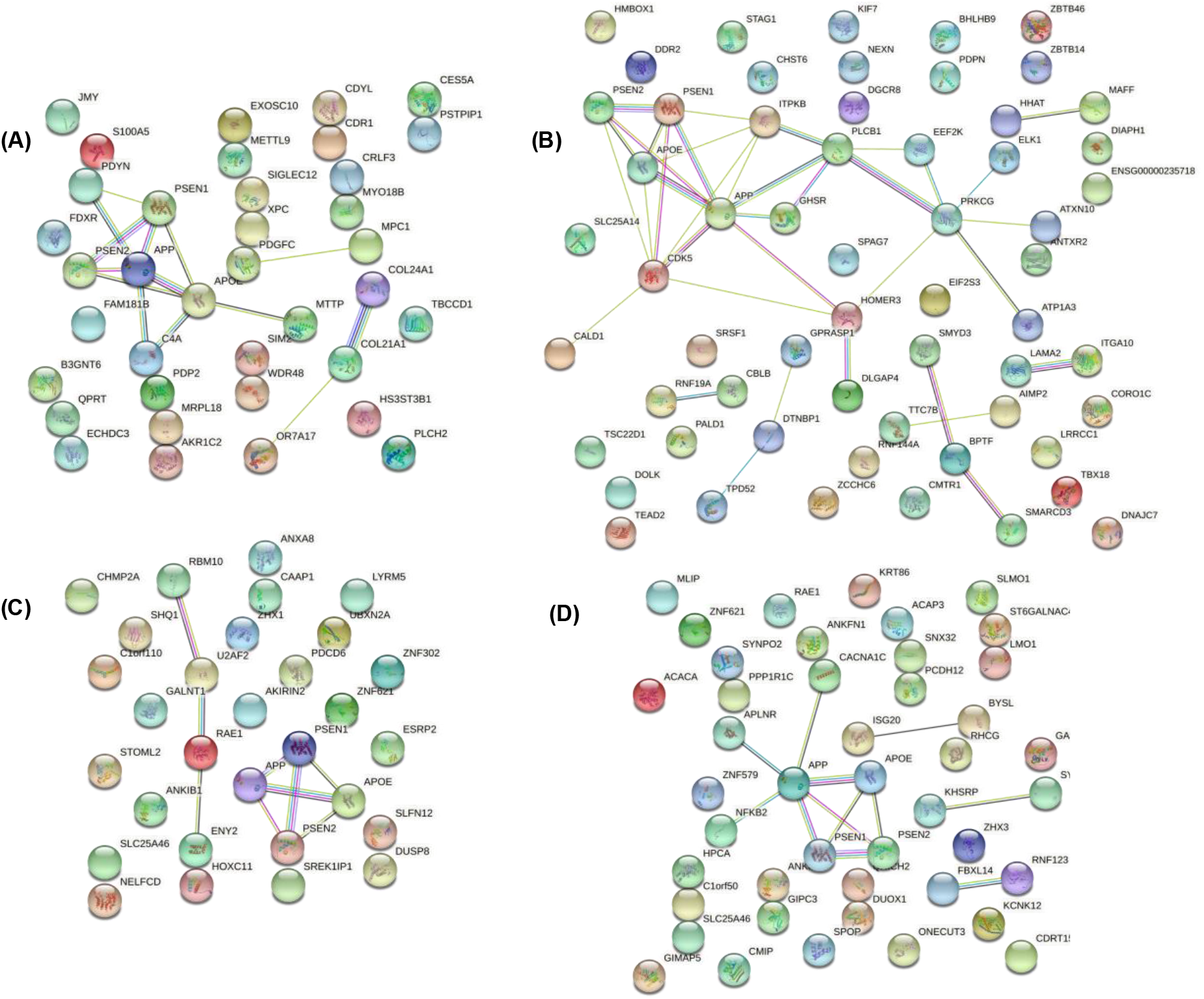
Protein-protein interaction (PPI) networks of the gene biomarkers for **(A)** Prefrontal cortex, **(B)** Middle Temporal Gyrus, **(C)** Hippocampus and **(D)** Entorhinal Cortex. The coloured nodes represent the proteins with first shell of interactions whereas the white nodes represent second shell of interactions. The proteins whose 3D structure are not known is shown by empty nodes. The coloured edges represent protein-protein interactions^40^.

## 5. Conclusion

The use of comprehensive machine learning models to identify potential gene biomarkers for Alzheimer’s disease is a significant step to determine the early treatment of AD patients. In this work, we propose a simple and robust framework to identify biologically important genes in the context of AD. There are three crucial aspects that corroborate the strength of the framework, (i) To identify the potential genetic markers of AD, probing the gene expression data from different brain tissue is more effective than analysing the combined profiles of expression level from all the regions together. In addition to that, incorporating a large sample size augments the credibility of the findings. (ii) The use of the best configured benchmark machine learning based feature selection model (wrapper approach) provided the most explaining gene subsets with the highest AD predictive power. (iii) To explain the biological significance, a strong validation is a must. Alongside conducting an extensive literature survey, the biological relevance is elucidated quantitatively by testing the biological significance of the obtained gene for two independent brain regions (Visual Cortex and Cerebellum). By employing the gene expression data of diseased vs. normal patients for four different brain regions to identify the biomarkers and incorporating them, our study has achieved, by far the highest prediction accuracy through optimally configured classification models.

In summary, we found several potential biomarkers, some of which are previously linked to AD such as ECHDC3, ZNF621, STOML2, TSC22D1, SIM2, CDK5, C4A, GHSR, PLCB1, ITPKB, NFKB2, CACNA1C, etc. and some novel biomarkers such as CORO1C, SLC25A46, RAE1, ANKIB1 CRLF3, PDYN, AK057435, and BC037880. Future work requires clinical and experimental testing of these gene candidates to identify potential prognostic biomarkers that can support the early diagnosis of Alzheimer’s disease or can be targeted at the gene level to prevent the disease. We will also extend the application of the proposed paradigm to discover novel potential markers for other complex diseases in future.

## Supplementary

## Machine Learning Model descriptions

### Random Forest (RF)

Given a training dataset, 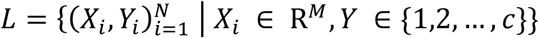, where *X*_*i*_ represents the variables or the feature set and *Y* denotes the corresponding label (class response variable). The number of training samples and features are denoted as *N* and *M* respectively. The random forest model (RF) is delineated below. For a given input *X*, let the prediction of the tree *T*_*k*_ is denoted by 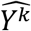. The random forest amalgamating *K* trees have the prediction given as:

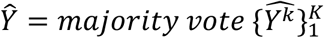

**Algorithm [1].**
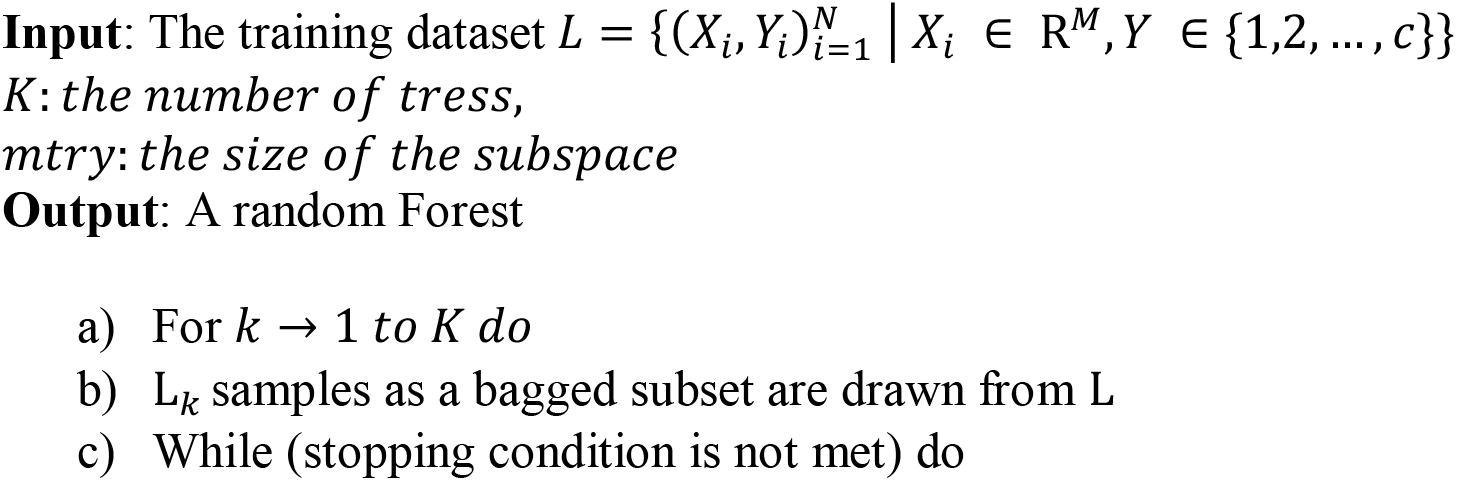
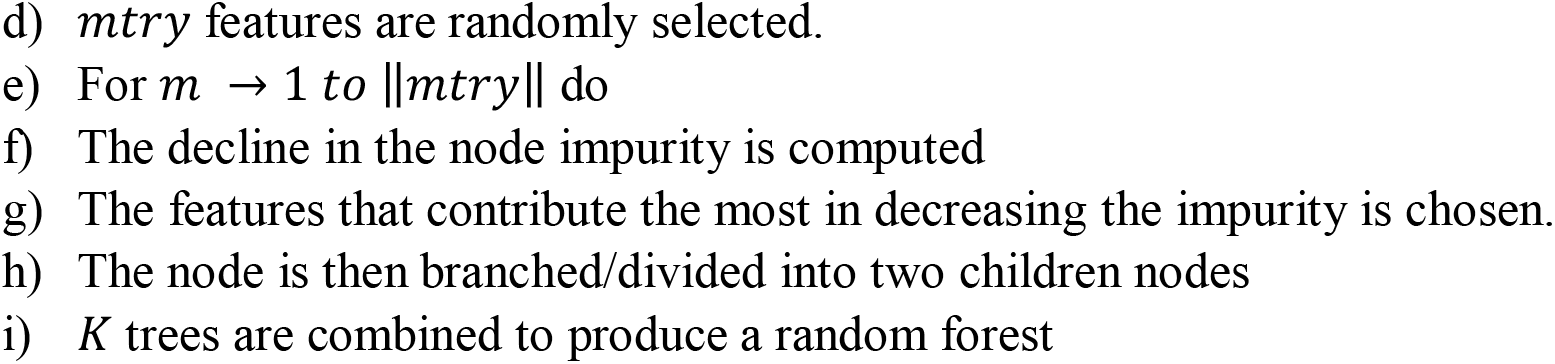

As the trees are grown from a bagged sample set, only a proportion of samples were leveraged to grow the tree also called *in-bag* samples. A small proportion of instance that is left out is called *out-of-bag* (OOB) samples that are employed to estimate the rate of prediction error called OOB error rate.

The OOB predicted value is given as: 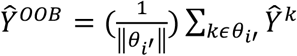, where 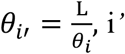 and *i* denotes the out-of-bag and in-bag sampled instances, ∥θ_*i*′_∥ is the cardinality/size of OOB instances, and the OOB prediction error is

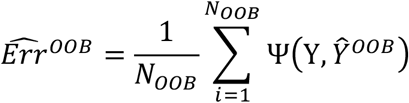

Here Ψ(.) is the error function and *N*_*OOB*_ is OOB sample’s size.

### Support Vector Machine (SVM)

An SVM classifier identifies and maximizes the most optimal hyperplane that separates the data points of each type of label (category). In a simple SVM model, the optimal hyperplane is evaluated on the basis of the distance between the support vectors [8]–[10] Once the hyperplane is evaluated using train data points, SVM allocates the new instances to a class based on its relative nearness from the trained data points [11]. For a given set of data points (*x*_*i*_, *y*_*i*_), *i* = 1,2, …., *m* where *x* ϵ R^*n*^, *y* ϵ **R**. Given a set of weight ***w***, The optimal hyperplane H is:

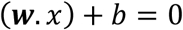

SVM classifier follows the constraints:

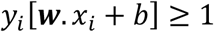

The optimization problem to minimize ***w*** (or maximize 2/∥***w***∥) is solved using a Lagrange function eq3:

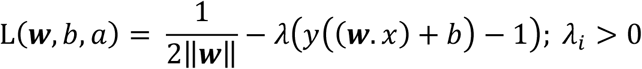

Here the λ is a Lagrange multiplier. Solving the partial derivatives for w and b to 0, the optimal hyperplane is built as:

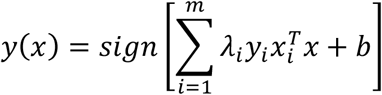

Several genomics studies have employed variations of SVM models as a classifier and retained excellent performance [12][13].

### Multiplicity Problem

For microarray dataset, different wrapper feature selection models identify a varying set of candidate genes as the important signature based on the prediction accuracy attained by the gene subset [2][3] This leads to the problem of multiplicity, especially for the case when the motivation is not only the prediction but also the identification of biologically relevant gene signatures[4][5]. This variation or the lack of uniqueness could be reasoned as different in the patient batch, differing analysis and varying technologies. Studies indicate that the difference in the gene subset is strongly influenced by the cohort that have been used for gene selection[3]. This problem has also been elaborated and discussed extensively in recent studies that too indicated the extremely small ratio of samples to genes in the microarray dataset is the most likely cause of this problem[6][7]. Unfortunately, this issue casts a false sense of trust in the results obtained by most studies falling under this paradigm of gene identification through wrapper approach. Subscribing to the notion of the studies investigating the problem of multiplicity, we lend credence to the combined set of genes that were obtained by both the methods (varSelRF and LASSO); and exclusively probed the biological significance of the common and repeatedly selected gene candidates.

---

**Figure S1.**
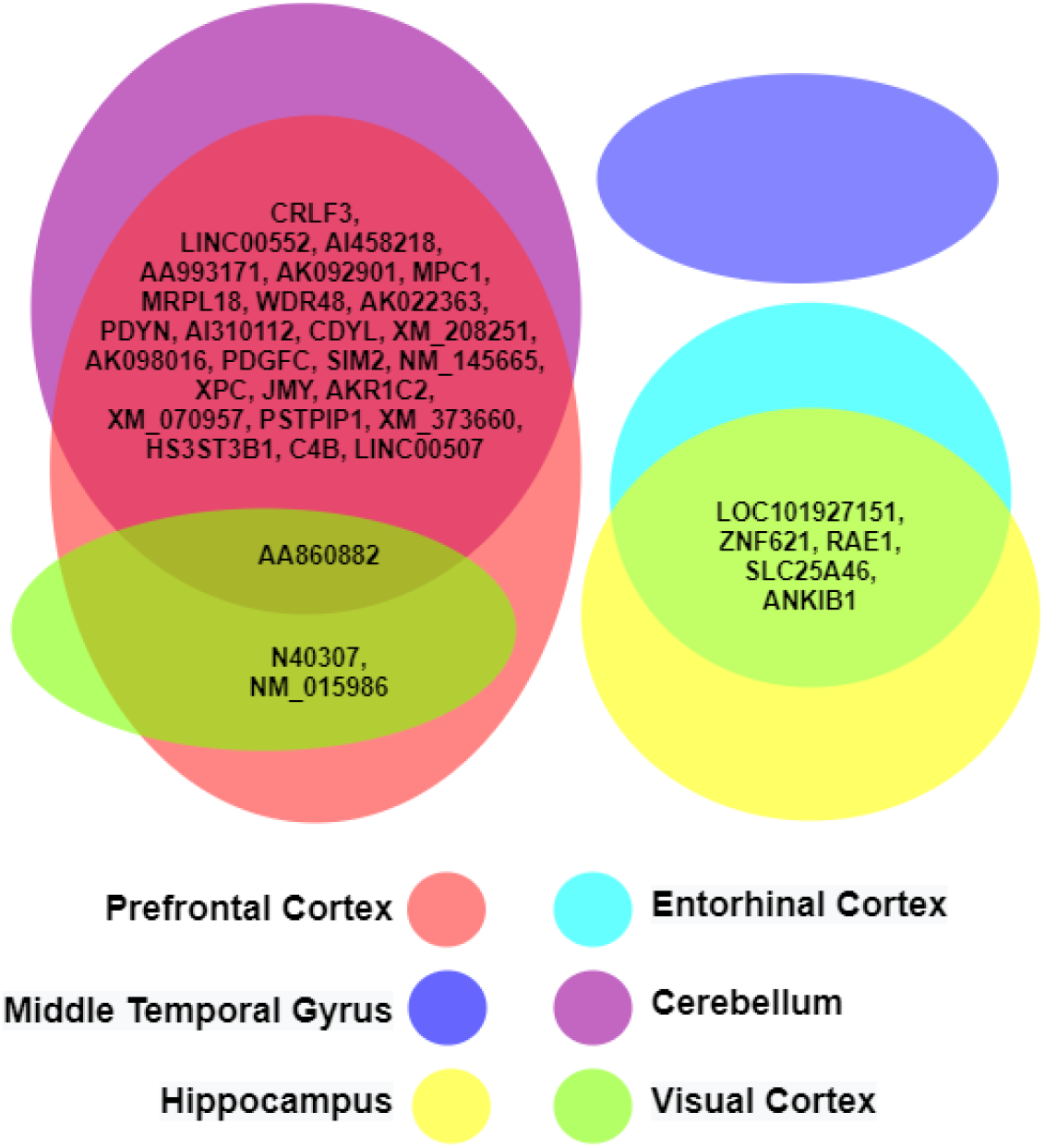
The common genes identified by the machine learning models within the different brain regions.

**Figure S2.**
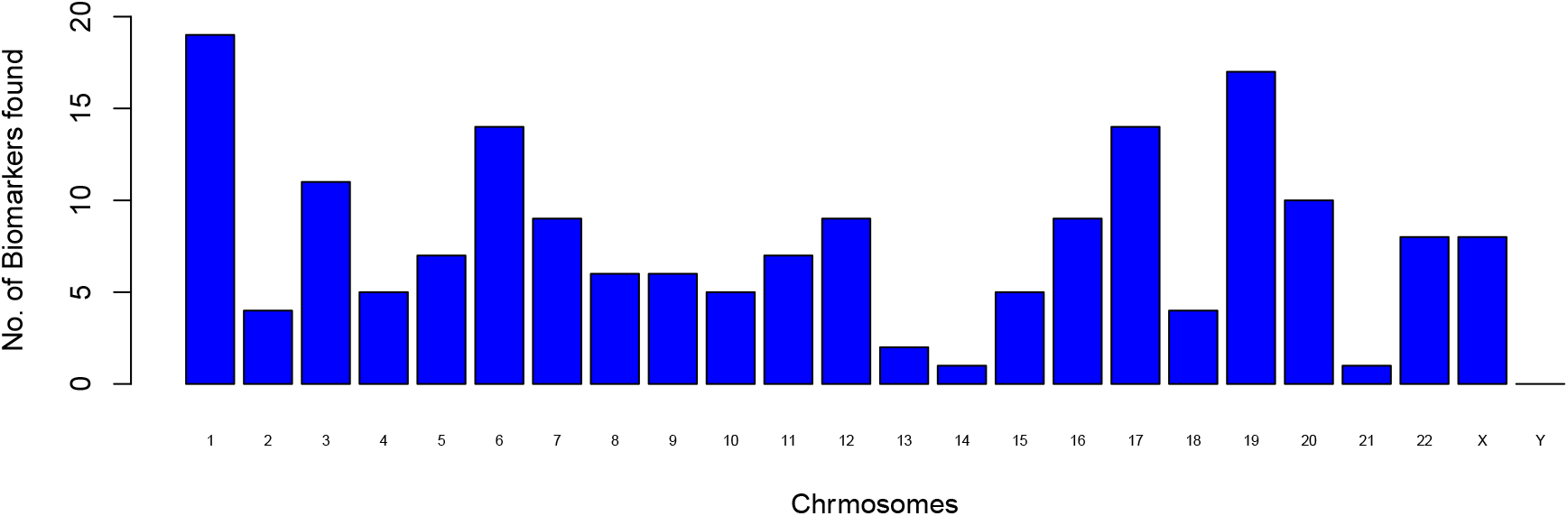
The number of biomarkers found in all the human chromosomes by our models.

**Figure S3.**
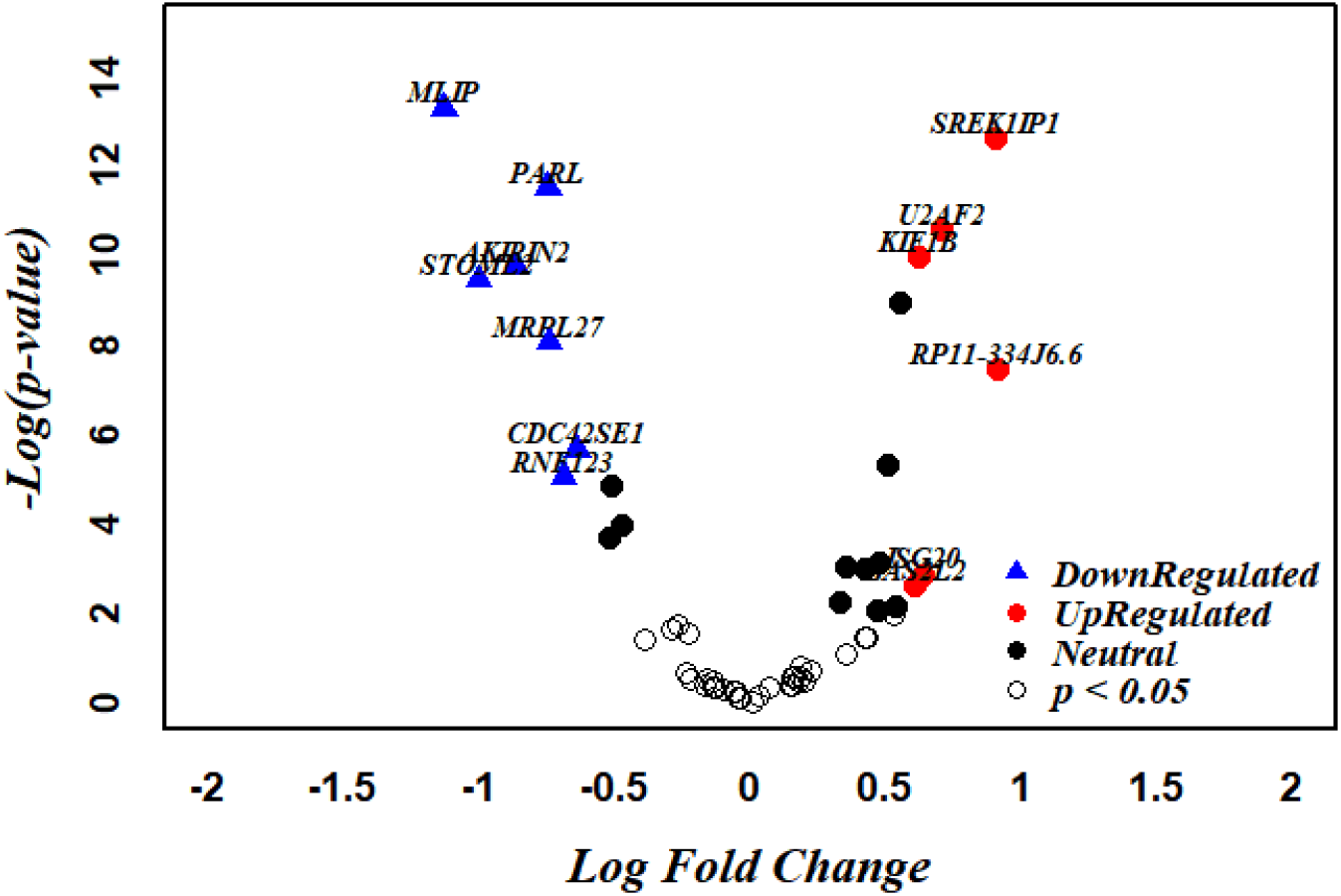
The up and down regulated biomarkers in the AD samples with respect to the normal ones identified from the GSE5281 expression data.

**Figure S4.**
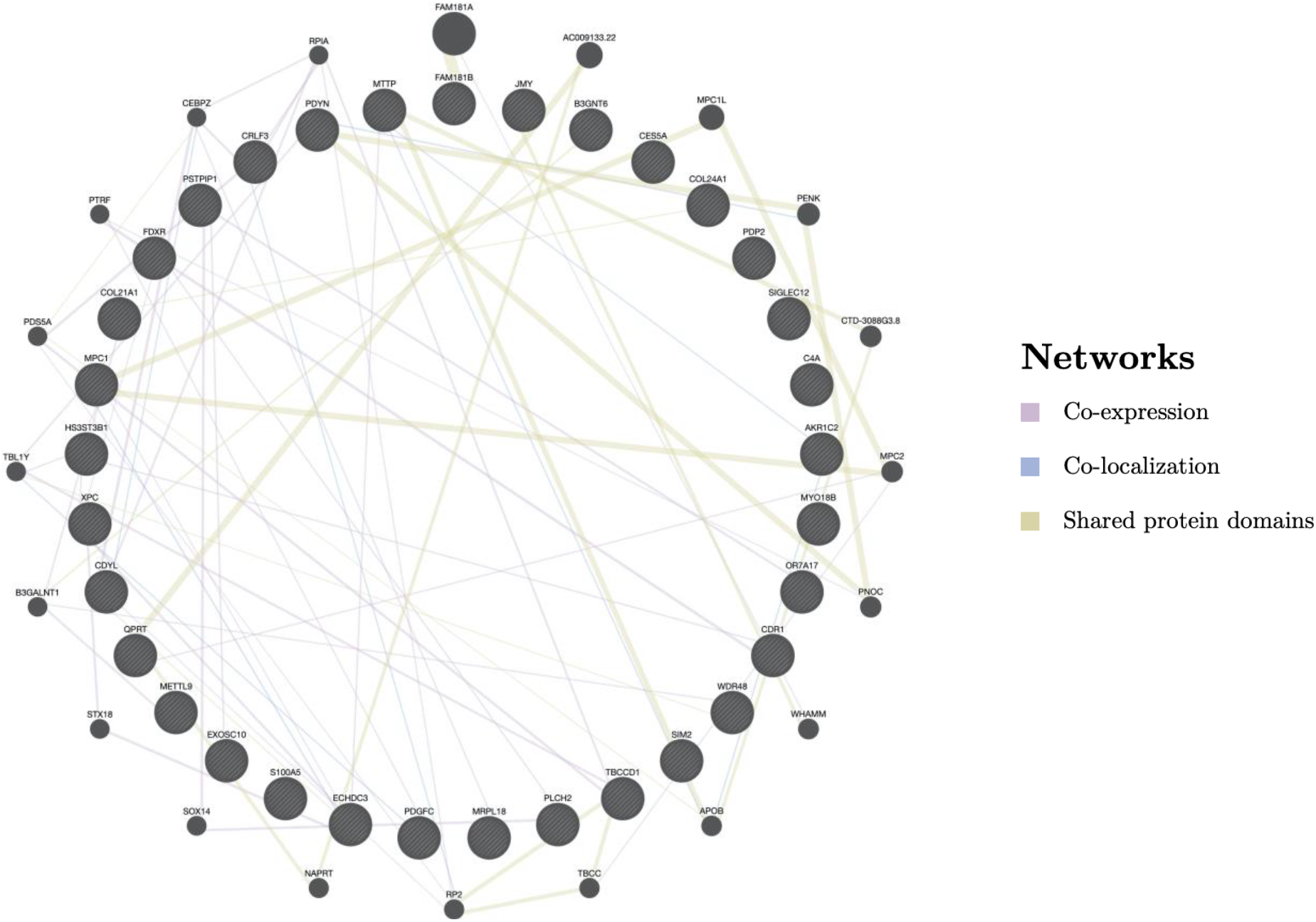
The GeneMania network analysis for all the biomarkers of PFC brain region. The biomarkers are shown by the stripped black circles. The solid-coloured lines represent the type of interactions in the biomarkers.

**Figure S5.**
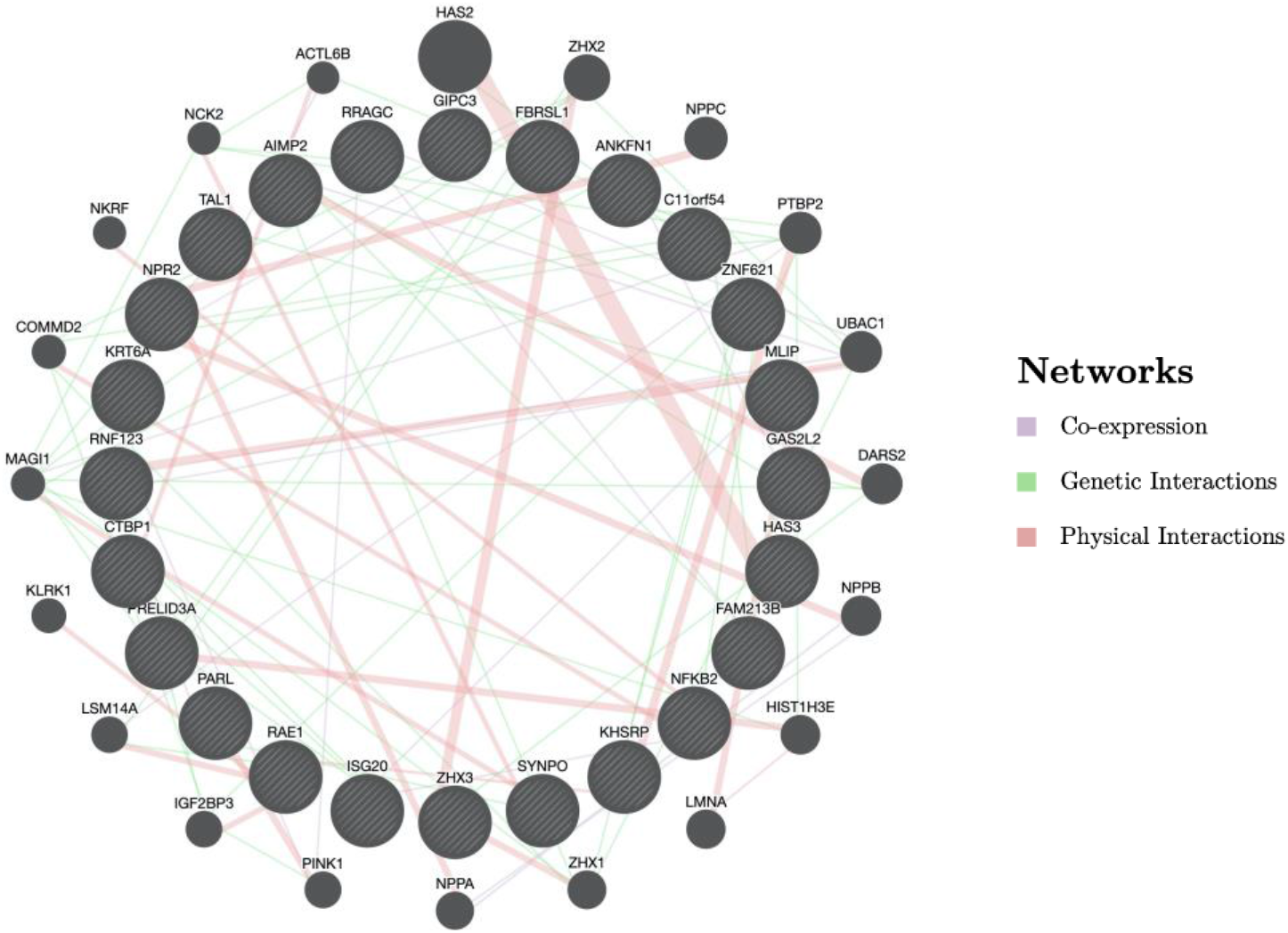
The GeneMania network analysis for all the biomarkers of EC brain region. The biomarkers are shown by the stripped black circles. The solid-coloured lines represent the type of interactions found within the biomarkers.

**Figure S6.**
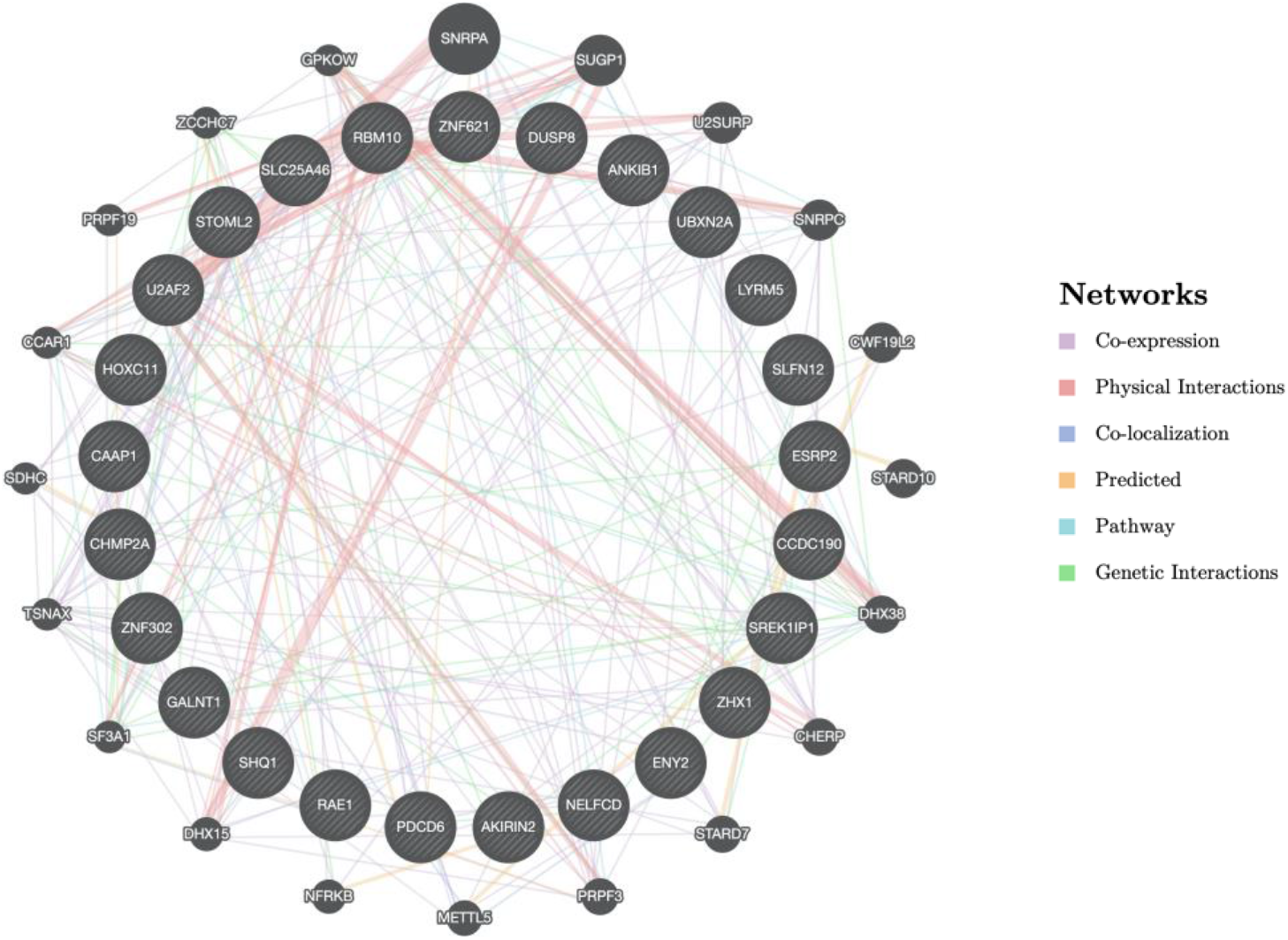
The GeneMania network analysis for all the biomarkers of H brain region. The biomarkers are shown by the stripped black circles. The solid-coloured lines represent the type of interactions found in the biomarkers.

**Figure S7.**
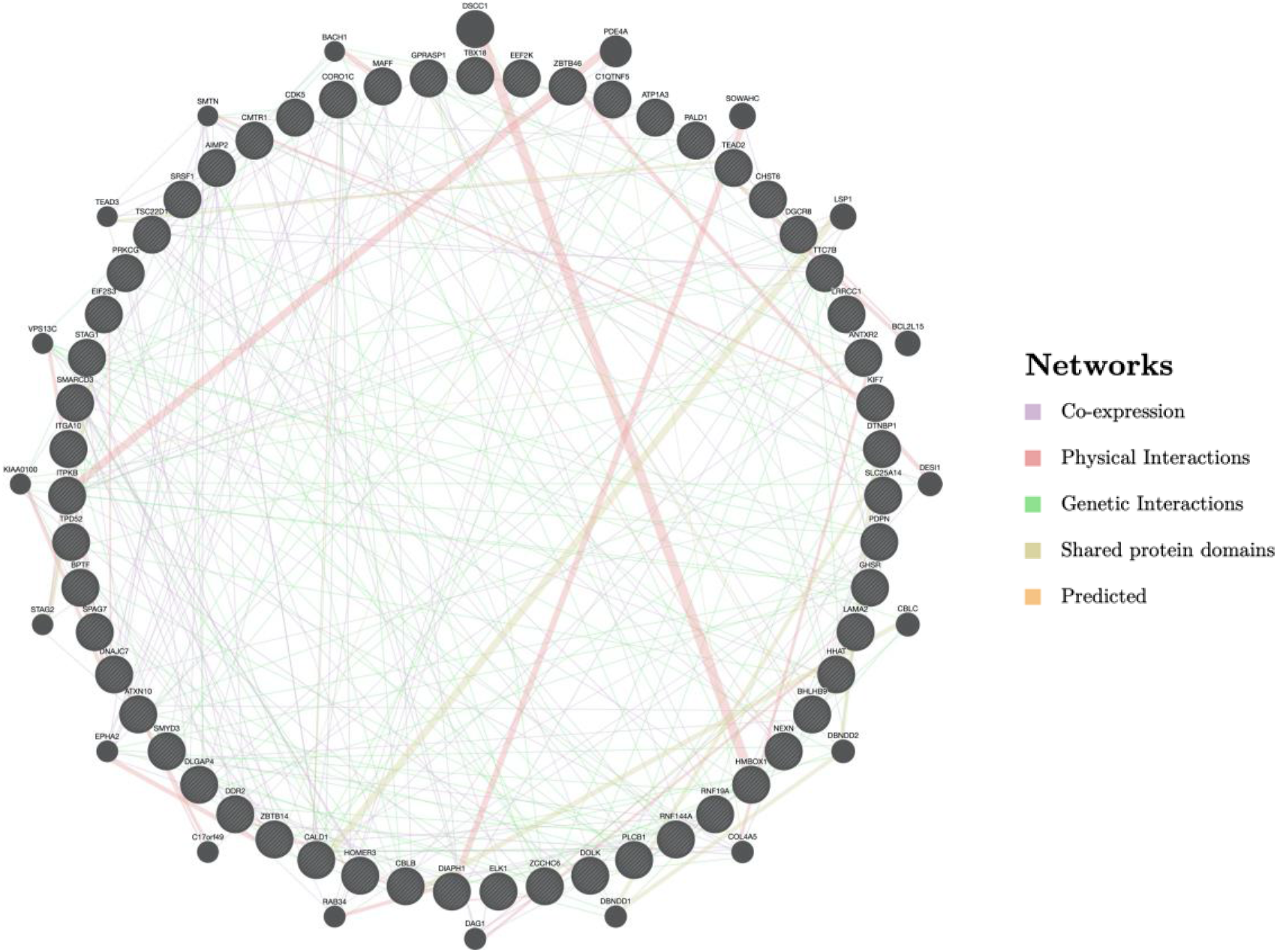
The GeneMania network analysis for all the biomarkers of MTG brain region. The biomarkers are shown by the stripped black circles. The solid-coloured lines represent the type of interactions found in the biomarkers.

**Table S1:** Summary of experimental designs and measurements of the gene expression datasets used in our study.

| Dataset | Dataset |
| --- | --- |
| GSE5281 | The signal value calculated by MAS 5 or GCOS software. |
| GSE48350 | GC-RMA normalized expression values |
| GSE4757 | The signal value calculated by MAS 5 or GCOS software. |
| GSE28146 | MAS5-calculated Signal intensity |
| GSE118553 | Normalized signal |
| GSE132903 | Normalized (log2 scale) with the lumiExpresso function (R-package Lumi) |
| GSE33000 | Normalized log10 ratio (Cy5/Cy3) representing test/reference |
| GSE44770 | Normalized log10 ratio (Cy5/Cy3) representing test/reference |
| GSE44771 | Normalized log10 ratio (Cy5/Cy3) representing test/reference |
| GSE44768 | Normalized log10 ratio (Cy5/Cy3) representing test/reference |

**Table S2:** The tuned hyperparameter sets for the different models used in our machine learning workflow.

| Model | Critical Parameters | Note |
| --- | --- | --- |
| varSelRF | ntree = 5000,<br>ntreeIterat = 2000,<br>vars.drop.frac = 0.2 | After tuning the default values remained the best value |
| LASSO | method = "glmnet",<br>lambda= seq(0.0001, 1, length = 5) | The alpha is not declared, setting 0 by default thus performing LASSO |
| RandomForest | ntree=500,<br>mtry= max( (number of gene) / 3 , 1) | ntree employed here is the default value which was tested against 250, 300 and 400. |
| Elastic Net | method = "glmnet",<br>alpha= seq(0,1,length=10),<br>lambda= seq(0.0001, 1, length = 5) | Both alpha and beta were searched within the given possible set |
| SVM | kernel= "radial",<br>degree = 7 | Degree of 7 is compared against 2 to 6 and remained the best suiting for the profiles |

**Table S3:** Alzheimer’s disease classification performance of the PFC biomarkers on VC and CR gene expression datasets.

| 5Fold CV | Visual Cortex |  |  |  |  | Cerebellum |  |  |  |  |
| --- | --- | --- | --- | --- | --- | --- | --- | --- | --- | --- |
|  | Acc | Sen | Spe | Pre | Mcc | Acc | Sen | Spe | Pre | Mcc |
| **Fold1_SVM** | 0.92 | 0.91 | 0.94 | 0.95 | 0.84 | 0.87 | 0.75 | 1.00 | 1.00 | 0.77 |
| **Fold1_RF** | 0.97 | 1.00 | 0.94 | 0.96 | 0.95 | 0.79 | 0.80 | 0.79 | 0.80 | 0.59 |
| **Fold1_EN** | 0.97 | 1.00 | 0.94 | 0.96 | 0.95 | 0.92 | 0.85 | 1.00 | 1.00 | 0.86 |
| **Fold2_SVM** | 0.97 | 1.00 | 0.94 | 0.96 | 0.95 | 0.92 | 0.95 | 0.88 | 0.91 | 0.84 |
| **Fold2_RF** | 0.92 | 0.95 | 0.88 | 0.91 | 0.84 | 0.90 | 0.91 | 0.88 | 0.91 | 0.79 |
| **Fold2_EN** | 0.97 | 0.95 | 1.00 | 1.00 | 0.95 | 0.97 | 0.95 | 1.00 | 1.00 | 0.95 |
| **Fold3_SVM** | 0.95 | 0.91 | 1.00 | 1.00 | 0.90 | 0.90 | 0.94 | 0.86 | 0.85 | 0.80 |
| **Fold3_RF** | 0.95 | 0.91 | 1.00 | 1.00 | 0.90 | 0.79 | 0.89 | 0.71 | 0.73 | 0.61 |
| **Fold3_EN** | 0.95 | 0.91 | 1.00 | 1.00 | 0.90 | 0.92 | 0.89 | 0.95 | 0.94 | 0.85 |
| **Fold4_SVM** | 0.85 | 0.87 | 0.81 | 0.87 | 0.68 | 0.90 | 0.90 | 0.89 | 0.90 | 0.79 |
| **Fold4_RF** | 0.85 | 0.87 | 0.81 | 0.87 | 0.68 | 0.85 | 0.86 | 0.83 | 0.86 | 0.69 |
| **Fold4_EN** | 0.85 | 0.87 | 0.81 | 0.87 | 0.68 | 0.92 | 0.95 | 0.89 | 0.91 | 0.85 |
| **Fold5_SVM** | 0.87 | 0.94 | 0.81 | 0.81 | 0.75 | 0.85 | 0.85 | 0.84 | 0.85 | 0.69 |
| **Fold5_RF** | 0.90 | 0.94 | 0.86 | 0.85 | 0.80 | 0.82 | 0.90 | 0.74 | 0.78 | 0.65 |
| **Fold5_EN** | 0.87 | 0.94 | 0.81 | 0.81 | 0.75 | 0.79 | 0.80 | 0.79 | 0.80 | 0.59 |
| **Avg_SVM** | 0.91 | 0.93 | 0.90 | 0.92 | 0.83 | 0.89 | 0.88 | 0.89 | 0.90 | 0.78 |
| **Avg_RF** | 0.92 | 0.94 | 0.90 | 0.92 | 0.83 | 0.83 | 0.87 | 0.79 | 0.82 | 0.66 |
| **Avg_EN** | 0.92 | 0.94 | 0.91 | 0.93 | 0.85 | 0.91 | 0.89 | 0.93 | 0.93 | 0.82 |

**Table S4:** The assessment metric obtained for VC and CR validating the AD prediction potential of the gene biomarker subset of PFC.

| Visual Cortex | Cerebellum |
| --- | --- |
| N40307, NM_015986, AA860882 | CRLF3, LINC00552, AI458218, AA993171, AK092901, MPC1, MRPL18, WDR48, AK022363, PDYN, AI310112, CDYL, XM_208251, AK098016, PDGFC, SIM2, NM_145665, XPC, JMY, AKR1C2, AA860882, XM_070957, PSTPIP1, XM_373660, HS3ST3B1, C4B, LINC00507 |
| **Common Marker** | CRLF3, AA860882 |

**Table S5:**
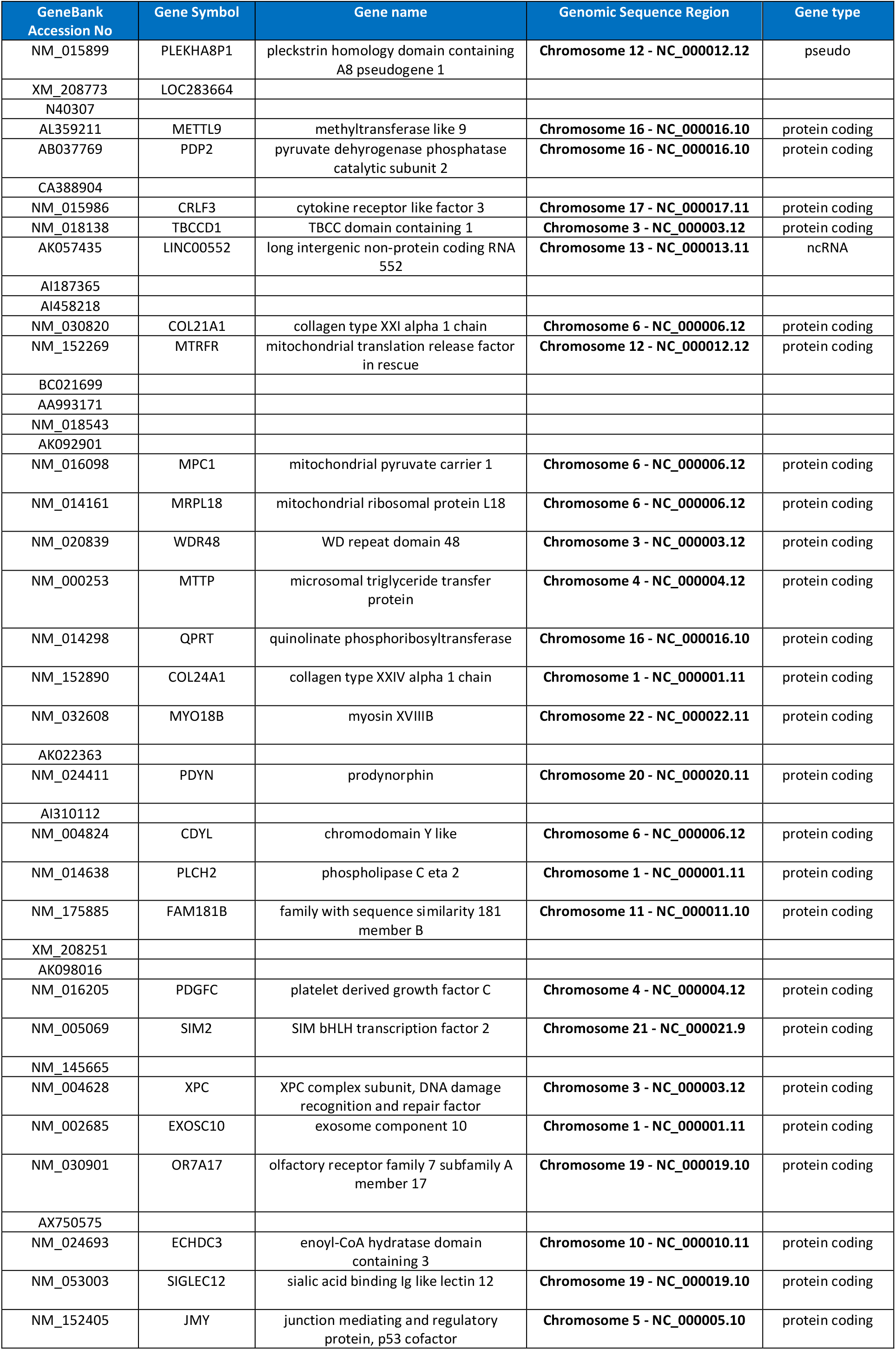
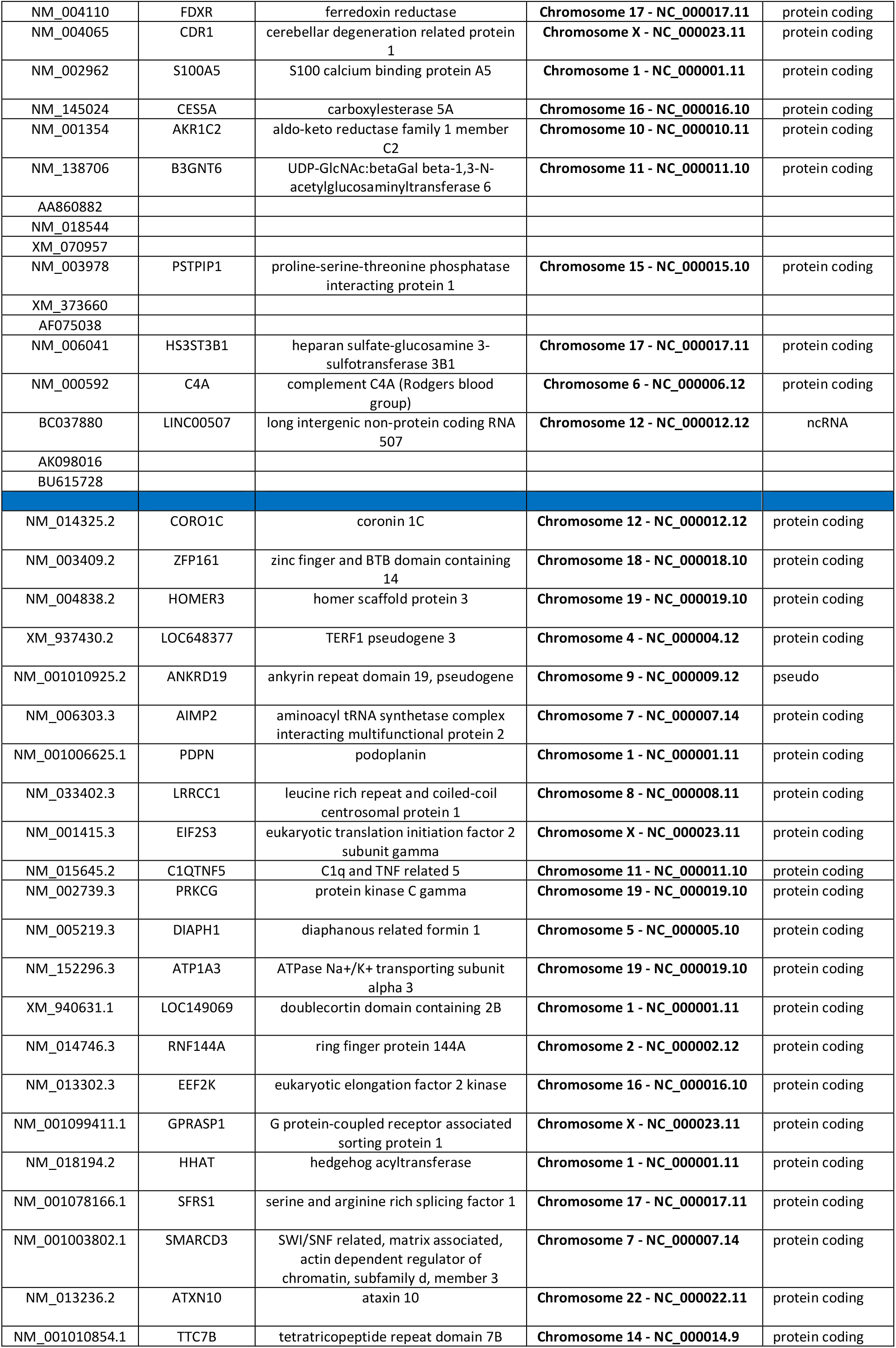
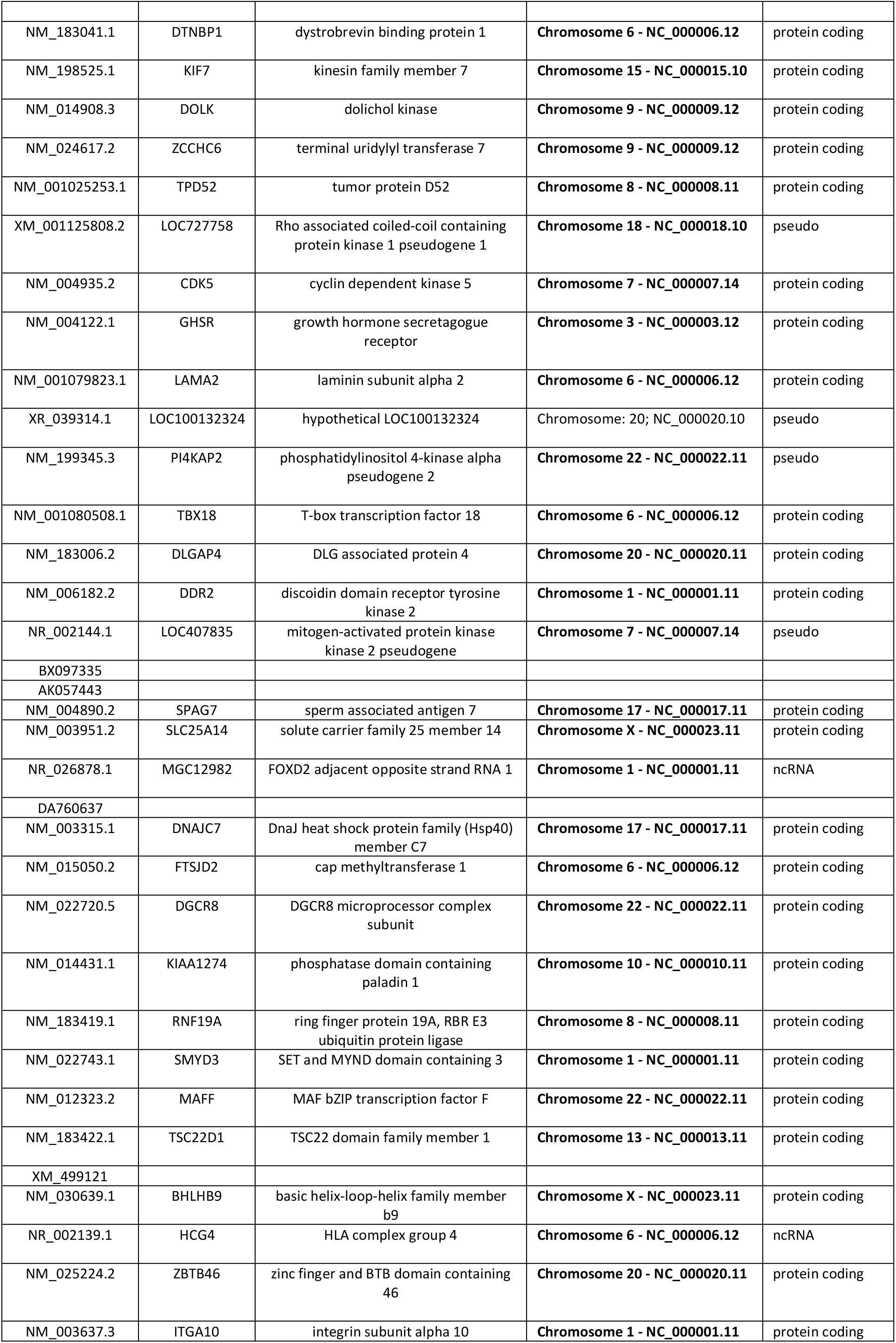
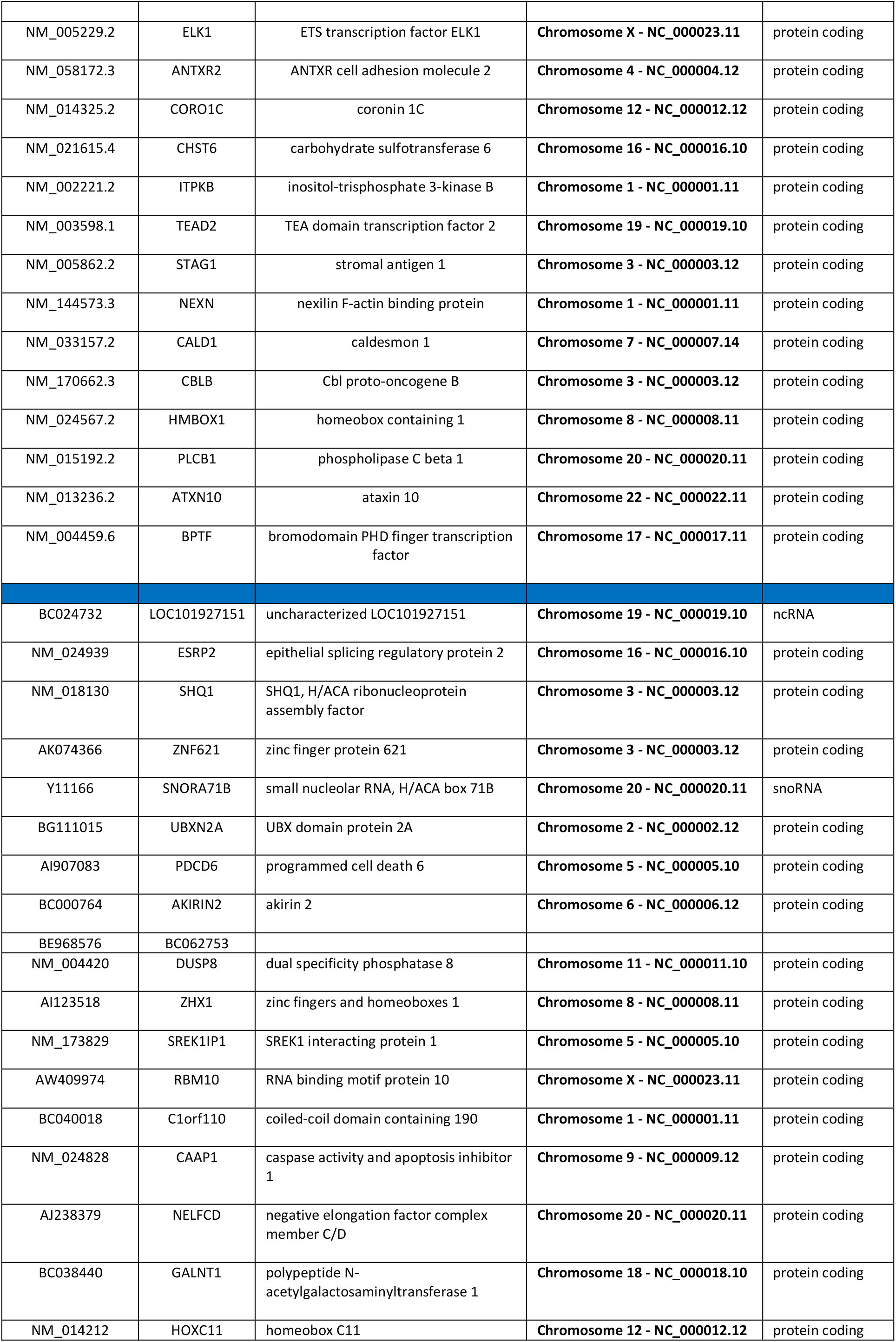
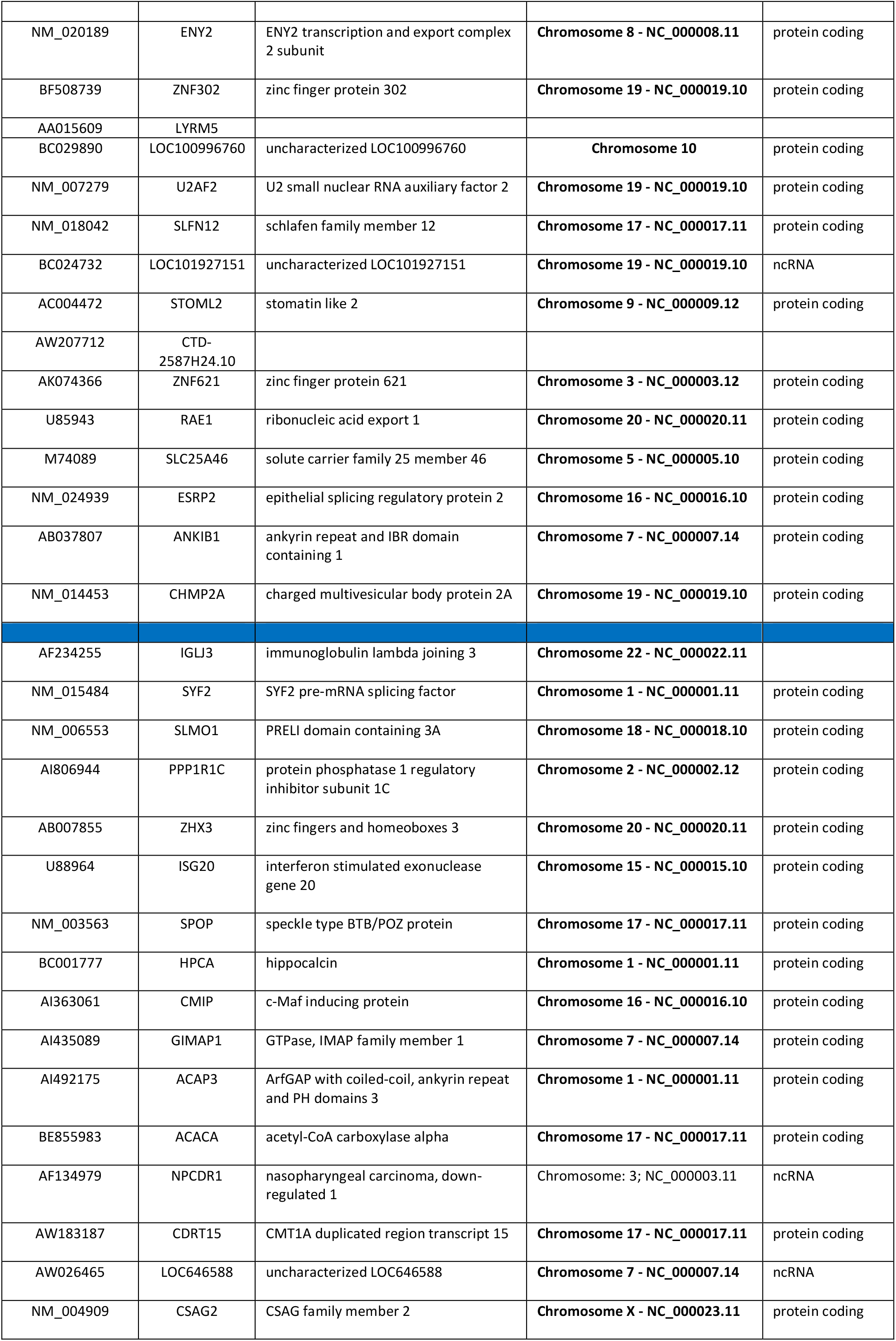
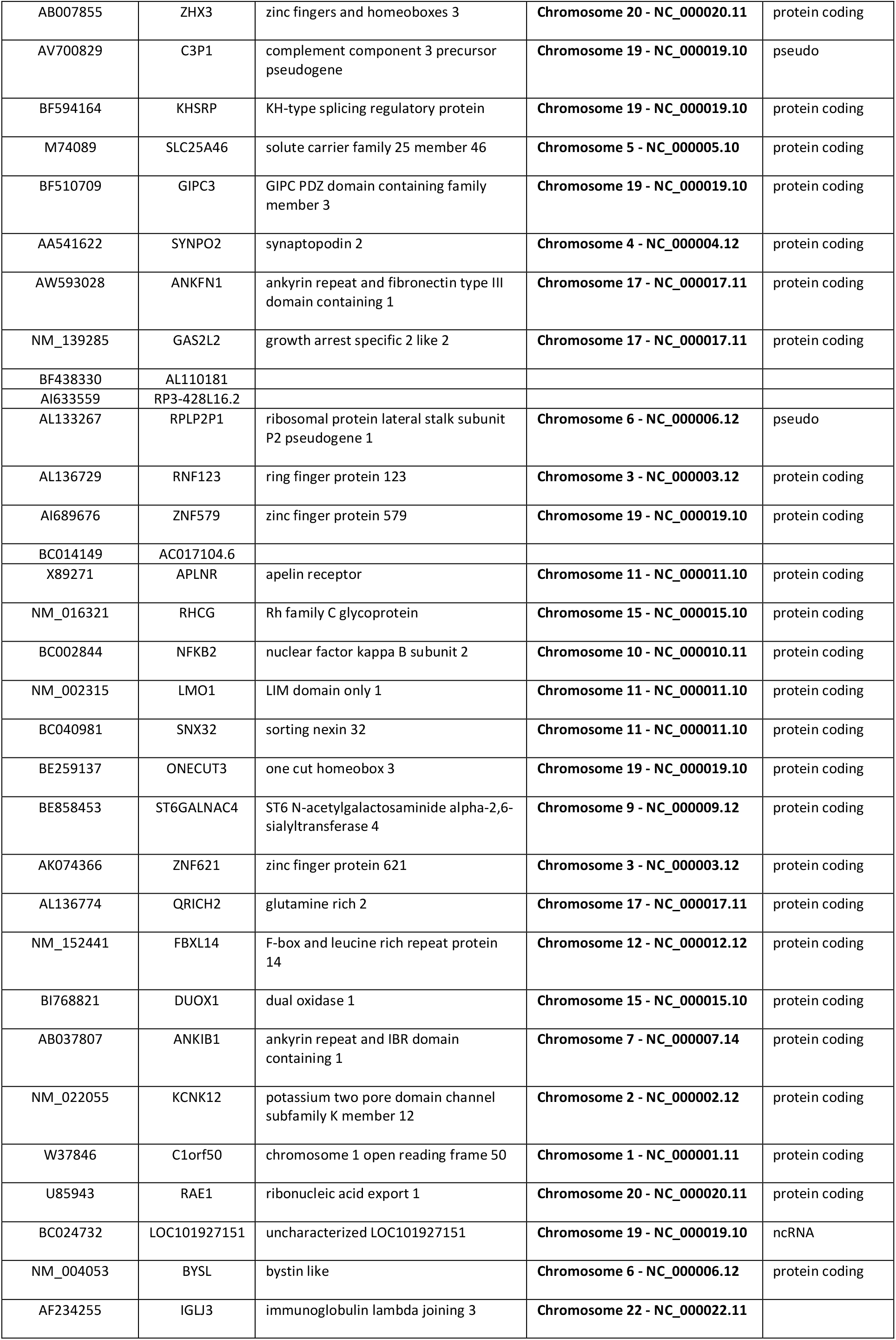
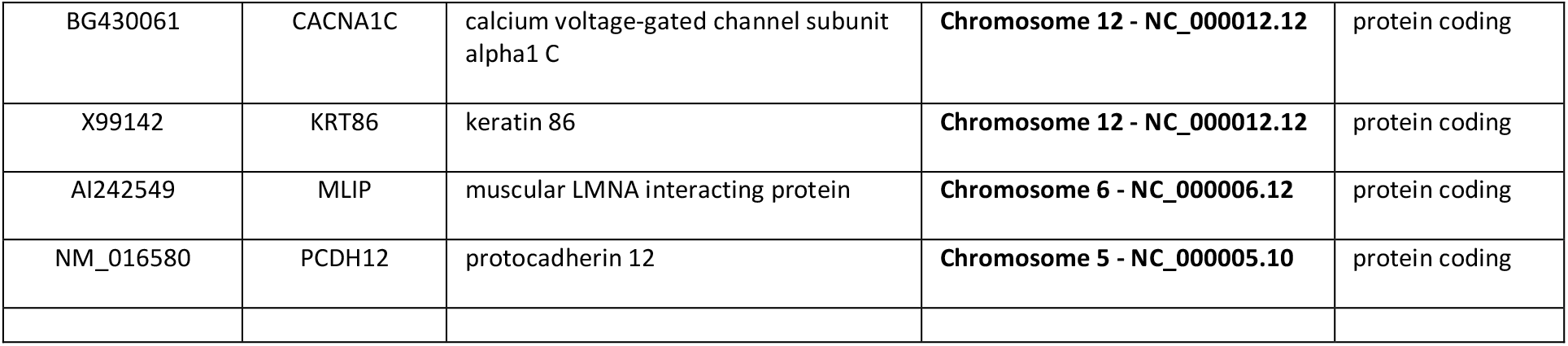
Bioinformatics analysis of the biomarkers found by the machine learning models.

